# Neural crest cells regulate optic cup morphogenesis by promoting extracellular matrix assembly

**DOI:** 10.1101/374470

**Authors:** Chase D. Bryan, Rebecca L. Pfeiffer, Bryan W. Jones, Kristen M. Kwan

## Abstract

The interactions between an organ and its surrounding environment are critical in regulating its development. In vertebrates, neural crest and mesodermal mesenchymal cells have been observed close to the eye during development, and mutations affecting this periocular mesenchyme can cause defects in early eye development, yet the underlying mechanism has been unknown. Here, using timelapse microscopy and four-dimensional cell tracking in zebrafish, we establish that genetic loss of neural crest impairs cell movements within the optic vesicle. At the ultrastructural level, neural crest cells are required for basement membrane formation specifically around the retinal pigment epithelium. Neural crest cells express the extracellular matrix crosslinking protein nidogen and, strikingly, ectopically expressing nidogen in the absence of neural crest partially restores optic cup morphogenesis. These results demonstrate that the neural crest is required for local establishment of ocular extracellular matrix superstructure, which in turn drives optic cup morphogenesis.

## Introduction

Vertebrate eye development begins with specification of the eye field, followed by a series of tissue movements that together comprise optic cup morphogenesis (Hilfer, 1983; Schmitt and Dowling, 1994; Schook, 1980; Walls, 1942). Initially, a pair of optic vesicles evaginate bilaterally from the developing forebrain; the bilayered optic vesicles give rise to the neural retina and retinal pigment epithelium (RPE). As these vesicles elongate, the connection between the vesicle and brain is constricted, generating the optic stalk. Multiple cell and tissue movements occur during invagination, the final stage of optic cup morphogenesis: the optic vesicles buckle and take on a hemispherical shape to enwrap the lens as it invaginates from the overlying ectoderm. Additionally, the optic fissure forms along the ventral side of the optic vesicle and optic stalk. Lineage tracing and live imaging experiments performed in zebrafish have enabled cellular-level analysis of the morphogenetic movements that occur during these stages of eye development. These experiments have revealed that during invagination, a subset of cells that arise on the medial layer of the optic vesicle migrate around the rim of the vesicle and eventually take up residence within the lateral layer, contributing to the neural retina; those which remain on the medial layer flatten and comprise the RPE (Heermann et al., 2015; Kwan et al., 2012; Li et al., 2000; Picker et al., 2009; Sidhaye and Norden, 2017). Despite characterization of these cellular movements, the cellular and molecular mechanisms underlying many aspects of these processes are not well understood.

In addition to these complex tissue movements and rearrangements, the optic vesicle undergoes morphogenesis in a complex environment containing multiple extraocular tissues, including the overlying ectoderm from which the lens will develop, the prospective brain, and the periocular mesenchyme (POM). Mesenchymal cells influence the development and morphogenesis of many epithelial organs, such as tooth and salivary gland, by producing growth factors and signaling molecules such as BMPs, WNTs, and FGFs (Thesleff, 2003; Wells et al., 2013), or through modifications to the extracellular matrix (ECM) that surrounds developing epithelia. For example, mesenchymal cells can influence morphogenesis through cleavage and destruction of ECM via proteolysis: in the developing mouse lung, mesenchymally-expressed matrix metalloproteinase-2 (MMP-2) is required for branching morphogenesis (Kheradmand et al., 2002). Conversely, mesenchymal cells can also promote epithelial morphogenesis through deposition of new ECM components such as laminin and nidogen, reviewed in (Nelson and Larsen, 2015).

Previous work has indicated a role for epithelial-mesenchymal interactions during optic cup morphogenesis, although the exact role and molecular nature of these interactions are poorly understood. The POM is a heterogeneous cell population in close proximity to the optic cup, comprised of neural crest cells and mesodermally-derived mesenchyme (Johnston et al., 1979), and multiple tissues in the mature eye are derived in part from these mesenchymal cell populations (Williams and Bohnsack, 2015). Recent work also indicates a role for POM in closure of the optic fissure, a developmental step subsequent to optic cup formation (Gestri et al., 2018). Yet in addition to these later roles, disruptions to transcription factors expressed in the POM during early eye development, such as *ap2a*, *pitx2* or *zic2*, lead to severe optic cup malformations (Bassett et al., 2010; Bohnsack et al., 2012; Gestri et al., 2009; Li and Cornell, 2007; Sedykh et al., 2017). These data suggest a critical role for the POM in regulating optic cup morphogenesis, possibly through regulating direct or indirect signaling to the optic cup.

Communication between these tissues appears to be bidirectional: the developing optic vesicle is known to signal to the migratory neural crest, in part through retinoic acid and PDGF signaling (Bohnsack et al., 2012; Eberhart et al., 2008; Lupo et al., 2011). In zebrafish *rx3* mutants, eye specification fails to occur; these embryos subsequently display aberrant craniofacial neural crest migration (Langenberg et al., 2008), indicating that the eye is at least partially responsible for facilitating proper neural crest migration. However, many of the molecules that mediate these bidirectional epithelial-mesenchymal interactions during optic cup morphogenesis have yet to be discovered.

Although there are hints that secreted morphogens such as Hedgehog or TGF-β may be crucial for crosstalk between the optic vesicle and the POM (Fuhrmann et al., 2000; Grocott et al., 2011; Sedykh et al., 2017), mesenchymal cells may also regulate optic cup development by modifying the ECM. A complex ECM has long been known to surround the developing optic vesicle throughout optic cup morphogenesis (Dong and Chung, 1991; Hendrix and Zwaan, 1975; Kwan, 2014; Peterson et al., 1995; Svoboda and O’Shea, 1987; Tuckett and Morriss-Kay, 1986), yet the dynamics of ECM deposition and remodeling around the developing eye are poorly understood. Only recently have specific roles of fibronectin (Hayes et al., 2012; Huang et al., 2011) and laminin (Bryan et al., 2016; Ivanovitch et al., 2013; Sidhaye and Norden, 2017) during optic cup morphogenesis been elucidated, and both of these ECM proteins are expressed by the optic vesicle itself. POM cells could modify the ocular ECM by expressing ECM-degrading proteins such as metalloproteases, or by providing structural ECM proteins such as nidogens: both are expressed by mesenchymal cells during morphogenesis of other organs. The roles of many ECM proteins during optic cup morphogenesis, especially those which may be produced by mesenchymal cells, have not been studied in detail, and their functions during this particular process remain a mystery.

The POM likely regulates multiple aspects of optic cup morphogenesis, yet many questions remain about the nature of the interactions between the POM and the developing eye. In this study, we sought to determine the role of the neural crest in regulating morphogenesis of the developing optic cup. When and where do neural crest cells interact with the optic vesicle during optic cup morphogenesis? Is the neural crest cell population actually required for optic cup morphogenesis? Are there morphogenetic events within the developing eye that depend on the neural crest? At a molecular level, how do neural crest cells interact with and regulate behaviors within the optic cup? Here we demonstrate that loss of neural crest cells, via two independent genetic methods, impairs optic cup morphogenesis. Using 4-dimensional timelapse imaging and computational methods, we pinpoint specific cell movements within the optic cup that are dependent on the neural crest. We further demonstrate that loss of neural crest leads to a dramatic disruption of basement membrane formation, but only around the RPE. Finally, we uncover a key molecular effector of neural crest crosstalk with the eye: our results indicate that neural crest cells regulate optic cup morphogenesis through deposition of nidogens, crucial modulators of ECM structure.

## Results

### Neural crest is in contact with the optic vesicle throughout optic cup morphogenesis

In zebrafish, optic cup morphogenesis occurs from 10-24 hours post fertilization (hpf), during which time the optic vesicles evaginate and undergo a series of stereotypical movements and shape changes to become the organized optic cup, comprised of the neural retina, retinal pigment epithelium (RPE), and lens. To begin to determine the nature of how the neural crest cell population might affect optic cup morphogenesis, we first sought to identify when and where neural crest cells interact with developing eye tissues. To visualize both the developing eye and neural crest, we crossed two transgenic zebrafish lines: *Tg(bactin2:EGFP-CAAX)^z200^*, in which GFP ubiquitously labels cell membranes, and *Tg(sox10:memRFP)^vu234^* (Kirby et al., 2006), in which neural crest cell membranes are marked with membrane-bound RFP. Using embryos from this cross, we performed 4-dimensional timelapse imaging during optic cup morphogenesis, from 12.5 hpf-24.5 hpf (Fig 1, Movies S1, S2). At the start of our imaging at 12.5 hpf, neural crest cells are coming into contact with the posterior margin of the optic vesicle (Fig 1A A’,). Initially, neural crest cells migrate anteriorly in the space between the prospective brain and optic vesicle (Fig 1B, Movie S1); beginning around 16 hpf, the developing optic stalk is gradually enwrapped by neural crest cells (Fig 1B’, Movie S2). Neural crest cells also migrate laterally and ventrally to encompass the posterior and ventral sides of the optic cup, appearing to be in close contact with the developing eye. By 24.5 hpf, the neural crest has entered the optic fissure and migrates toward the space between the neural retina and lens (Fig 1D’, *arrow*). By the end of optic cup morphogenesis, neural crest-derived cells have encapsulated the RPE side of the optic cup (Fig 1D).

**Figure 1.**
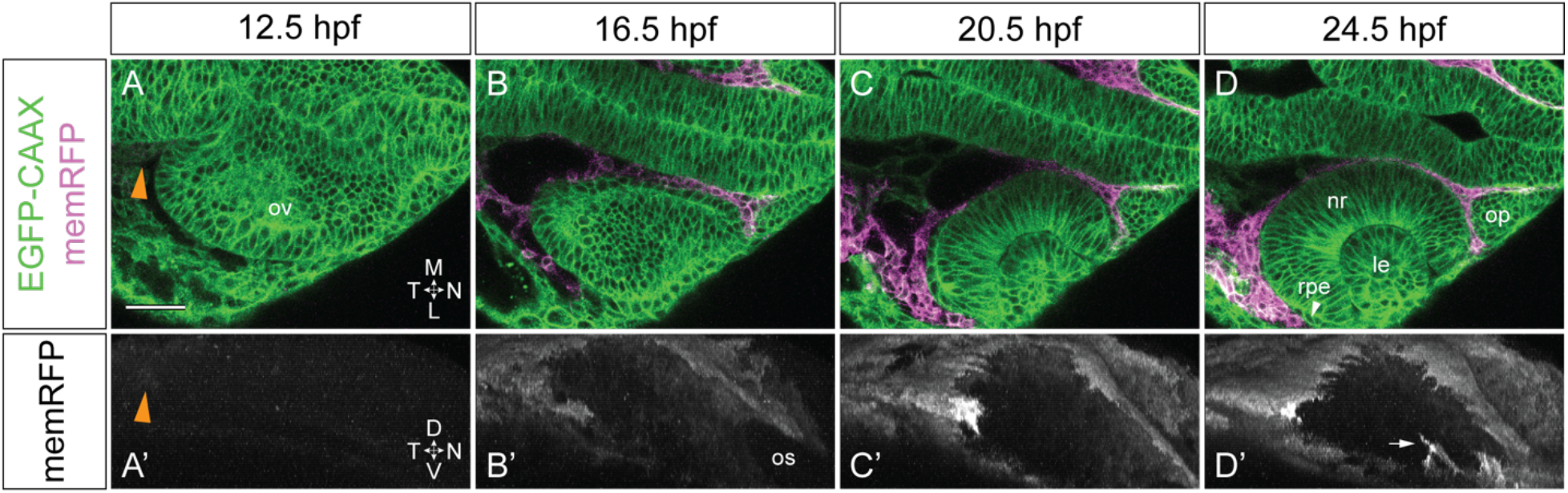
Neural crest is in contact with the optic vesicle throughout optic cup morphogenesis. Live imaging time series from 12.5-24.5 hpf of a *Tg(bactin2:EGFP-CAAX);Tg(sox10:memRFP)* double transgenic embryo. (A-D) Dorsal view, single confocal sections from a 4D dataset. (A’-D’) Lateral view, 3D rendering of the RFP channel from the same dataset as shown in (A-D). Orange arrowheads in A, A’ indicate neural crest cells beginning to express memRFP. White arrowhead in D indicates retinal pigment epithelium between neural crest and neural retina. White arrow in D’ indicates neural crest-derived cells entering the optic fissure. Scale bar, 50 μm. ΔT between z-stacks, 3.5 minutes. ov, optic vesicle; os, optic stalk; nr, neural retina; rpe, retinal pigment epithelium; le, lens; op, olfactory placode. M, medial; L, lateral; D, dorsal; V, ventral; N, nasal; T, temporal.

### Neural crest cells are required for proper optic cup morphogenesis

Multiple studies have suggested that there is developmental crosstalk between the developing eye and the neural crest (Bohnsack et al., 2012; Eberhart et al., 2008; Grocott et al., 2011; Langenberg et al., 2008; Sedykh et al., 2017). Since we observed neural crest cells in contact with the optic vesicles at early stages of eye development, we sought to determine whether neural crest cells are required for proper optic cup morphogenesis. To test the requirement for neural crest, we used two independent genetic models, both of which exhibit a widespread depletion of neural crest.

The zebrafish *tfap2a;foxd3* double mutant has been demonstrated to exhibit a strong loss of neural crest cells (Arduini et al., 2009; Wang et al., 2011). We crossed adult *tfap2a^ts213^;foxd3^zdf10^* heterozygote carriers to two transgenic lines: *Tg(bactin2:EGFP-CAAX)* and *Tg(sox10:memRFP)*. Crossing these two transgenic/double heterozygote lines enabled us to visualize the optic cup as well as to assay the presence of any remaining neural crest cells in the *tfap2a;foxd3* double mutant. At 24 hpf, *tfap2a;foxd3* double mutants display mild but reproducible optic cup morphogenesis defects. At the dorsal-ventral midpoint of the optic cup, the nasal side of the neural retina is flatter than in sibling control embryos and fails to completely enwrap the lens, indicative of a defect in optic cup invagination (Fig 2A, B). Quantification of optic cup invagination angle indicates a significant decrease in the extent to which the retina enwraps the lens in *tfap2a;foxd3* double mutants (37.8±1.9°) compared to wildtype controls (47.3±1.8°; Fig 2G). The optic fissure, a cleft-like structure along the ventral side of the optic stalk and optic cup, is also aberrant in *tfap2a;foxd3* double mutants. At 24 hpf, control embryos display two closely apposed fissure margins (Fig 2H) while the margins in *tfap2a;foxd3* double mutants are wider set, indicating that optic fissure development is abnormal (Fig 2I). Visualizing *sox10:memRFP*-positive cells indicates that, as expected, neural crest cells are substantially reduced in the vicinity of the optic cup in *tfap2a;foxd3* double mutants (Fig 2E, L) compared to wildtype controls (Fig 2D, K), with a notable absence on the nasal side of the optic cup (Fig 2L). Since incrosses of *tfap2a;foxd3* heterozygote adults yield both single as well as double mutant genotypes, we characterized *tfap2a* and *foxd3* single mutants as well. Neither single mutant displays as apparent optic cup morphogenesis defects or decrease in invagination angle as the double mutant (Fig S1A, B, E), likely due to the presence of more neural crest cells in either single mutant compared to the double mutant (Fig S1C, D).

**Figure 2.**
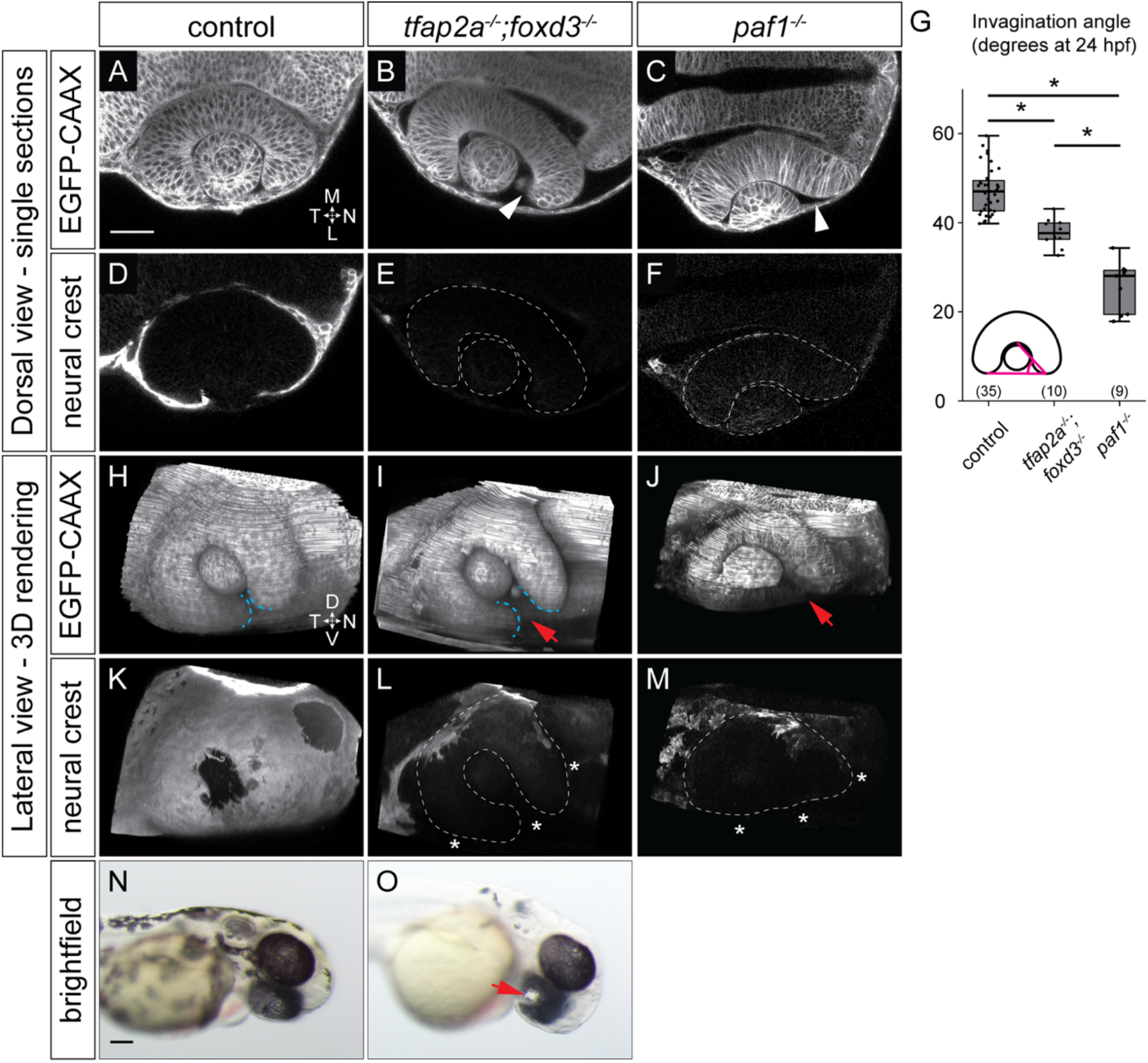
Optic cup morphogenesis is disrupted in neural crest mutants. (A-F) Dorsal view, single confocal sections of 24 hpf *Tg(sox10:memRFP)*-positive control (A, D), *tfap2a;foxd3* double mutant (B, E), and *paf1* mutant (C, F) embryos. Sections shown are at the dorsal/ventral midpoint of the lens. EGFP-CAAX was used to visualize optic cup morphology: (A, B) *Tg(bactin2:EGFP-CAAX)*; (C) injected with EGFP-CAAX mRNA. White dashed lines (E, F, L, M) show optic cup boundaries. White arrowheads (B, C) mark the nasal retina failing to fully enwrap the lens. (G) Quantification of invagination angles, measured as shown in inset diagram. ^*^P<0.001 using Welch’s t-test. (Control vs. *paf1* mutant, P=6.59×10- 7; control vs. *tfap2a;foxd3* double mutant, P=2.52×10-7; *tfap2a;foxd3* double mutant vs. *paf1* mutant, P=0.0001.) Results are from 2-3 experiments; n (embryos) shown at base of the graph. (H-M) Lateral view, 3D renderings of the embryos in (A-F). Blue dashed lines mark optic fissure margins. Note wideset optic fissure margins in the *tfap2a;foxd3* mutant in (I); no discernable margins are visible in the *paf1* mutant at 24 hpf. White asterisks (L, M) indicate regions missing neural crest. (N, O) Brightfield images of 52 hpf control and *tfap2a;foxd3* double mutant embryos. Red arrows (I, J, O) indicate coloboma. Scale bars: 50 μm in (A), 100 μm in (N). M, medial; L, lateral; D, dorsal; V, ventral; N, nasal; T, temporal.

As a second means of testing the requirement for neural crest cells in optic cup morphogenesis, we assayed optic cup morphogenesis in *alyron^z24^* (*paf1*) mutants in which neural crest development is severely impaired (Cretekos and Grunwald, 1999). Although ubiquitously expressed, disruptions to components of the RNA Polymerase II-Associated Factor (Paf1) complex result in severe reductions in neural crest gene expression, coupled with developmental defects in neural crest derived tissues (Akanuma et al., 2007; Langenbacher et al., 2011; Nguyen et al., 2010). We observe that optic cup invagination is more severely disrupted in *paf1* mutants (25.6±3.6°) when compared to wildtype controls or *tfap2a;foxd3* double mutants (Fig 2C, G). We also visualized neural crest in *paf1* mutants and saw a substantial reduction in *sox10:memRFP*-positive cells surrounding the optic cup at 24 hpf (Fig 2F, M), similar to the neural crest loss seen in the *tfap2a;foxd3* double mutant. However, as *paf1* and other members of the Paf1 complex are expressed ubiquitously (Nguyen et al., 2010; Thisse and Thisse, 2004), it is possible that *paf1* also plays an intrinsic role in development of the optic vesicle itself, which could account for the more severe morphogenesis defects we observe in the *paf1* mutants. Thus, further analysis on the role of neural crest in optic cup morphogenesis was carried out solely using the *tfap2a;foxd3* double mutant.

Previous studies have suggested a role for the periocular mesenchyme in closure of the optic fissure along the ventral side of the retina and optic stalk (Gestri et al., 2018; Hero, 1990; Hero et al., 1991; James et al., 2016; Lupo et al., 2011; Weiss et al., 2012). Consistent with these data, we see optic fissure defects and gaps in ocular pigmentation (indicative of coloboma) in 59.38% of *tfap2a;foxd3* double mutants at 52 hpf versus 7.62% of control embryos (n=19/32 and n=8/105, respectively; data from three separate clutches; Fig 2N, O). However, these previous studies have focused largely on later stages of optic fissure fusion, during which the POM appear to play an active role. Therefore, our observations at 24 hpf demonstrate that the neural crest additionally plays a role in the early stages of optic cup morphogenesis.

### TGF-β signaling is unaffected by loss of neural crest, while Pax2a expression is expanded

The finding that neural crest cells are required for early stages of optic cup morphogenesis raised the possibility that neural crest cells were providing a signaling cue to the developing optic cup; an intriguing candidate we first tested was TGF-β signaling. Work performed in chick optic vesicle explants demonstrated that POM cells are necessary for proper RPE specification and development, a requirement that could be bypassed with treatment with the TGF-β family member Activin (Fuhrmann et al., 2000). Other experiments have suggested that neural crest cells repress lens specification through TGF-β signaling to ensure proper positioning of the lens (Grocott et al., 2011). Thus, we sought to determine whether the neural crest is necessary for TGF-β signaling to the developing eye in zebrafish. Using an antibody against phospho-Smad3 to detect sites of active TGF-β signaling, we did not detect any differences in phospho-Smad3 localization between control and mutant optic cups at 24 hpf (Fig 3A, C vs 3B, D; Fig S2A-D). This result indicates that the neural crest subpopulation of POM is not required for proper TGF-β signaling at the end of zebrafish optic cup morphogenesis.

**Figure 3.**
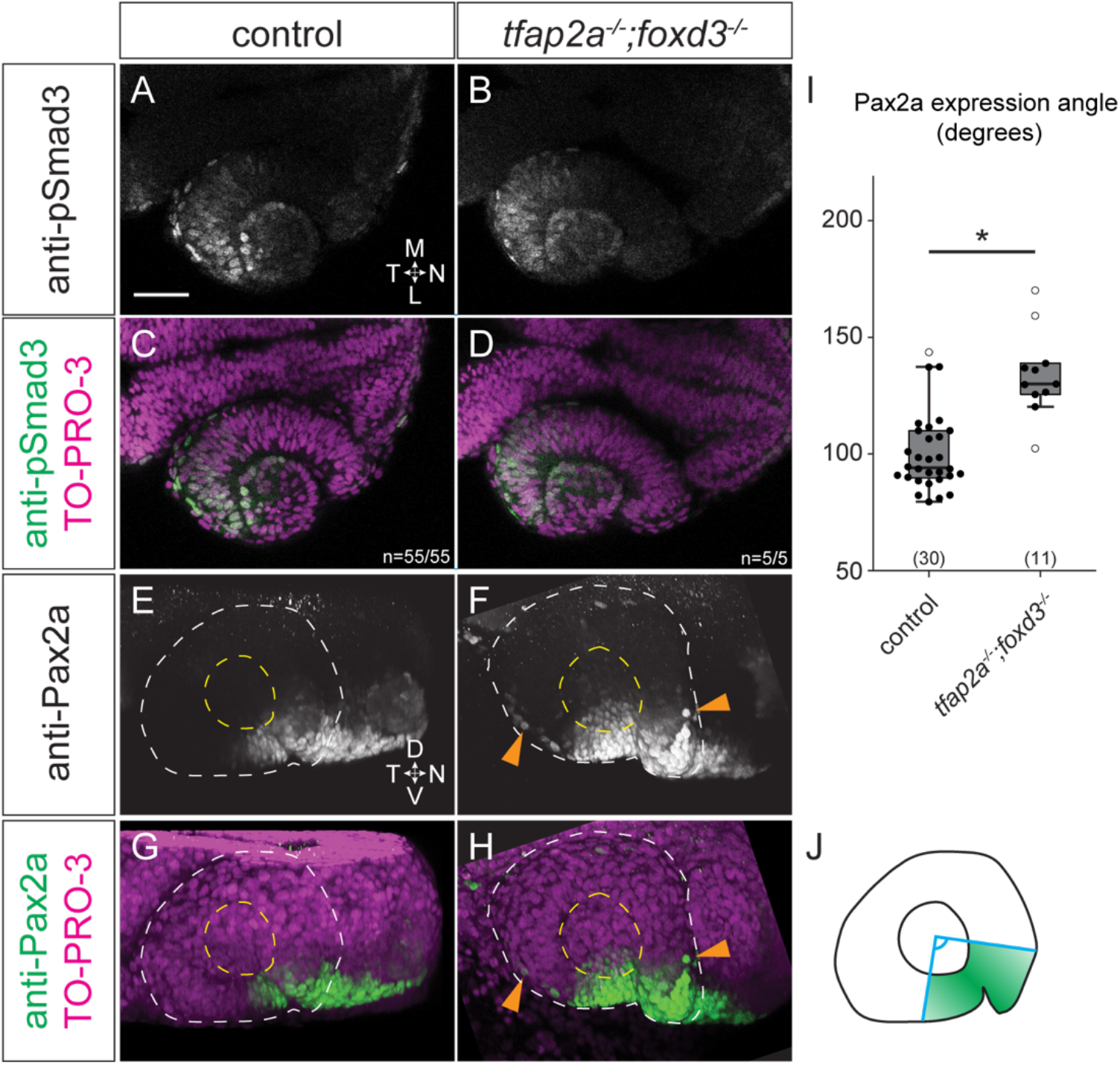
At 24 hpf, *tfap2a;foxd3* double mutants display normal TGF-beta signaling, while Pax2a expression expands into the RPE. (A-D) Dorsal view, single confocal sections of 24 hpf control (A, C) and *tfap2a;foxd3* double mutant (B,D) optic cups stained with anti-phospho-Smad3. Sections shown are at the dorsal/ventral lens midpoint. (E-H) Lateral view, 3D renderings of 24 hpf control (E, G) and *tfap2a;foxd3* double mutant (F,H) optic cups stained with anti-Pax2a. White dashed circles denote the boundary of the optic cup, yellow dashed circles display the boundary of the lens. Orange arrowheads in (F, H) indicate RPE cells which ectopically express Pax2a. (I) Angle measurements of the Pax2a expressing portion of control and *tfap2a;foxd3* double mutant optic cups. ^*^P=5.39×10^-5^ using Welch’s t-test, white circles are outliers. Results are from 3 experiments; n (embryos) shown in the base of the graph. Angles were measured from the lateral surface as diagrammed in (J), with the vertex of the angle set at the center of the lens. Nuclei were counterstained with TO-PRO-3 (magenta); merges shown in (C, D, G, H). M, medial; L, lateral; D, dorsal; V, ventral; N, nasal; T, temporal.

As we observed morphogenetic abnormalities in nasal and ventral portions of the optic cup in *tfap2a;foxd3* double mutants, we hypothesized that the neural crest might be required for some aspect of optic cup patterning. Using an antibody against Pax2a, a transcription factor expressed in the ventral optic cup and optic stalk (Fig 3E, G), we found that Pax2a expression is expanded into the RPE layer in *tfap2a;foxd3* double mutants, in cells located more dorsally or temporally than observed in wildtype eyes (arrowheads in Fig 3F, H). We quantified the portion of the optic cup which expressed Pax2a as an angle (schematized in Fig 3J) and find that expression is significantly expanded in the *tfap2a;foxd3* double mutant (134.2±10.5°) compared to control embryos (100.3±5.9°; Fig 3I). This expansion was consistently observed in *tfap2a;foxd3* double mutants as well as both *tfap2a* and *foxd3* single mutants (Fig S2E-H). In other models of ventral optic cup mispatterning, especially due to aberrant or increased Hedgehog signaling, Pax2a is expanded not just into the RPE layer but throughout the ventral hemisphere of the retina (Lee et al., 2008; Sedykh et al., 2017). This phenotype we observe here is distinct in its restriction to the RPE layer, and suggests that perhaps gross mispatterning of the ventral optic cup does not occur when neural crest is lost, but rather, some aspect of cell movements within the optic cup is disrupted.

### Neural crest cells are required for proper cell movements within the optic vesicle

We observed optic cup invagination defects and ectopic Pax2a expression in cells in the RPE layer at 24 hpf, which suggested that cell movements within the optic cup might be disrupted in the *tfap2a;foxd3* double mutant. We therefore sought to pinpoint when and where cell movements are disrupted in the *tfap2a;foxd3* double mutant. Determining how widespread movement defects are would provide clues regarding the nature and role of the interactions between the optic vesicle and neural crest. We directed our attention to two movements executed by cells which reside in the medial layer of the optic vesicle, as these are the cells potentially interacting with the neural crest, as visualized in Figure 1. The first movement is that of prospective RPE cells, while the second, rim involution, involves a subset of cells that migrate around the developing optic cup rim from the medial layer into the lateral layer, the prospective neural retina (Heermann et al., 2015; Kwan et al., 2012; Picker et al., 2009; Sidhaye and Norden, 2017). In mutant optic cups, expanded Pax2a expression in the RPE layer could result from failure of Pax2a-expressing cells to undergo rim movement into the neural retina, thus remaining in the apparent RPE layer of the optic cup; we hypothesized that rim involution might be disrupted in the absence of neural crest cells. To test this possibility, as well as to determine how widespread the cell movements are which depend on the neural crest, we performed live imaging and 4-dimensional cell tracking of optic cup morphogenesis in wildtype and *tfap2a;foxd3* double mutant embryos (Fig 4, Movie S3).

**Figure 4.**
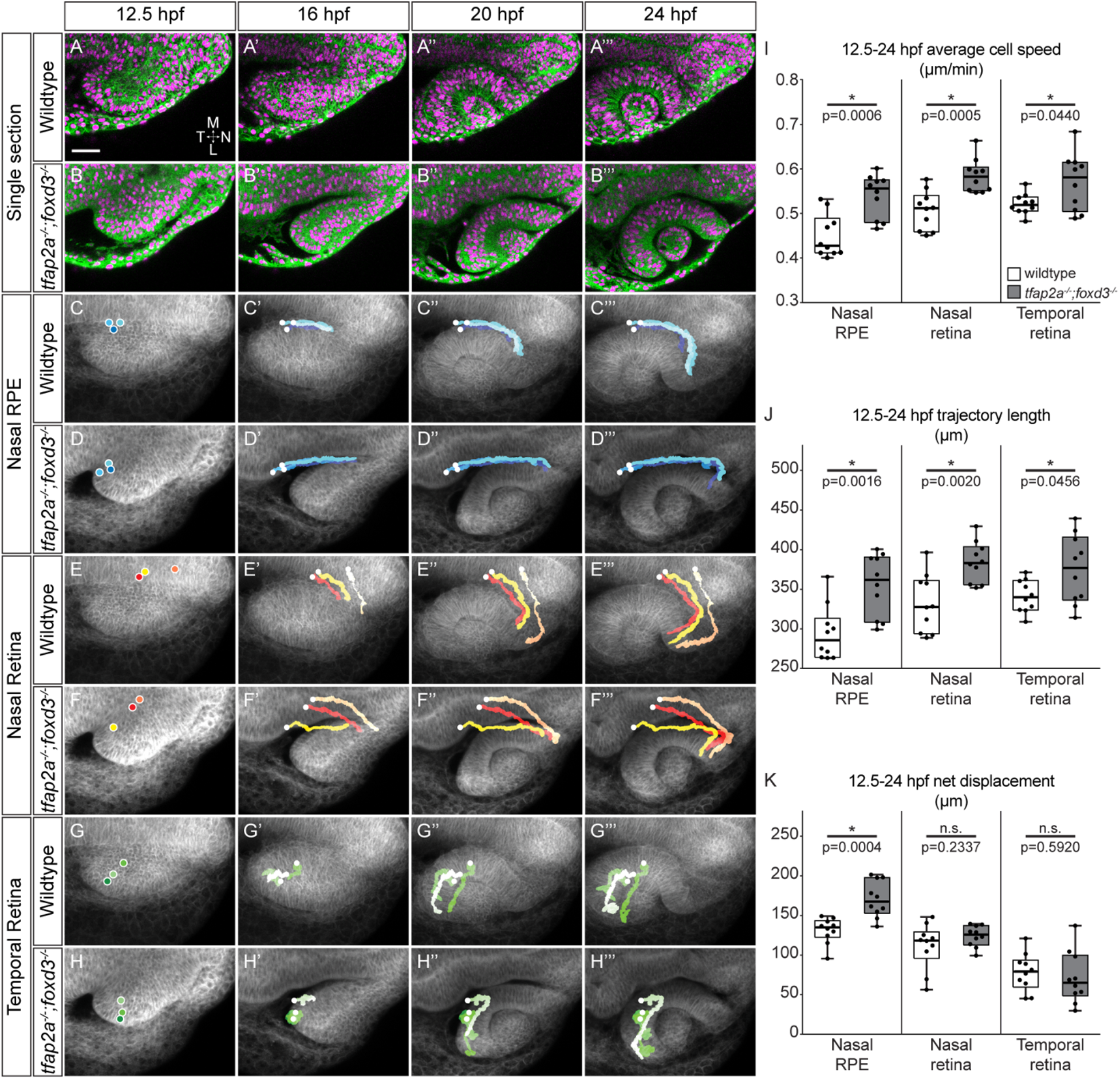
Cell movements throughout the optic cup are disrupted in *tfap2a;foxd3* double mutants. Live imaging time series of optic cup morphogenesis from 12.5-24 hpf of *Tg(bactin2:EGFP-CAAX)* wildtype and *tfap2a;foxd3* double mutant embryos. (A-B’’’) Dorsal view, single confocal sections from wildtype (A-A’’’) and *tfap2a;foxd3* double mutant (B-B’’’) 4D datasets. EGFP-CAAX (green) labels cell membranes, while H2A.F/Z-mCherry (magenta) labels nuclei. (C-H’’’) Average projections of membrane (EGFP-CAAX) channel through ~60 μm centered at the dorsal/ventral midpoint of the optic vesicle with indicated nuclear trajectories (nasal RPE, nasal retina, or temporal retina) overlaid. Trajectories were measured in 3D and generated by adding nuclear signal selections over time. (I-K) Average cell speed (I), total trajectory length (J), and overall net displacement (K) from cells which contribute to indicated region at 24 hpf. n=10 cells from each region (5 cells each from 2 embryos per genotype), backtracked from 24 to 12.5 hpf. P-values calculated using Welch’s t-test. Scale bar, 50 μm. ΔT between z-stacks, 2.75 minutes (wildtype) or 2.5 minutes (*tfap2a;foxd3*). M, medial; L, lateral; N, nasal; T, temporal.

As we saw distinct invagination defects in the nasal hemisphere of 24 hpf *tfap2a;foxd3* double mutant optic cups, we began by selecting nasal RPE and retina cells in their final positions at 24 hpf. We then tracked these cells retrospectively to establish their origins and movements from optic vesicle stage; a subset of tracked cells and trajectories is shown in Figure 4. First we visualized RPE cell movements and found clear changes in the trajectory of *tfap2a;foxd3* double mutant cells. While the nasal RPE population arises in the same starting position in both wildtype and double mutant embryos, wildtype nasal RPE cells execute a gradual, arc-like trajectory toward the nasal-lateral portion of the RPE and continually move toward the anterior and lateral sides of the embryo (Fig 4C-C’’’, Movie S4). In contrast, the *tfap2a;foxd3* nasal RPE cells move in a straight, anterior direction until 19.5 hpf, when they make a sudden, 90-degree turn in the lateral direction (Fig 4D-D’’’, Movie S5). To our surprise, when we quantify these cell movements (Fig 4I-K), we find that nasal RPE cells move faster and farther in *tfap2a;foxd3* double mutants (average speed = 0.57±0.02 μm/min; total distance = 385.75±15.71 μm) than wildtypes (average speed = 0.45±0.05 μm/min; total distance = 303.01±34.06 μm; Fig 4I, J). Additionally, net displacement of nasal RPE cells is increased in *tfap2a;foxd3* double mutants (170.76±14.55 μm) compared to wildtypes (131.17±10.16 μm), indicating that these cells do not arrive at the correct position within the optic cup. Taken together, these data confirm that neural crest is required for proper migration and positioning of nasal RPE cells during optic cup morphogenesis.

Through cell tracking, we also identified defects in the movements of nasal retinal cells. We find that both wildtype (Fig 4E-E’’’, Movie S6) and *tfap2a;foxd3* double mutant nasal retinal cells (Fig 4F-F’’’, Movie S7) arise in equivalent domains (Kwan et al., 2012) and follow a similar trajectory during optic cup morphogenesis, with one key difference: while wildtype retinal cells undergo a lateral turn away from the midline around 17.75 hpf (Movie S6), that same turn is delayed in the *tfap2a;foxd3* mutant until approximately 19.5 hpf (Movie S7). Similar to the nasal RPE population, nasal retinal cells move faster and farther in the *tfap2a;foxd3* double mutant (average speed = 0.60±0.04 μm/min; total distance = 402.80±17.33 μm) than wildtype (average speed = 0.53±0.03 μm/min; total distance = 352.41±29.99 μm; Fig 4I, J), although net displacement of these cells is unchanged (wildtype=111.55±18.18 μm, vs. *tfap2a;foxd3*=124.08±8.43 μm).

Our observation of expanded Pax2a expression into the temporal RPE suggested that cell movements may be disrupted on that side of the optic cup as well. Despite a lack of gross morphological defects on the temporal side of the optic cup, we found differences between wildtype and *tfap2a;foxd3* double mutant cell trajectories on this side of the optic vesicle as well. In wildtype embryos, the cells which undergo rim involution on the temporal side begin to migrate around the rim of the optic cup at approximately 20 hpf, with noticeable movement into the neural retina visible by 22 hpf (Fig 4G-G’’’, Movie S8). In the *tfap2a;foxd3* double mutant, these same cells do not begin migrating around the rim until 22 hpf (Fig 4H-H’’’, Movie S9). As observed with cells on the nasal side of the retina, temporal retinal cell speeds and distances traveled are significantly increased in the *tfap2a;foxd3* double mutant (average speed = 0.57±0.04 μm/min; total distance = 375.01±27.53 μm) compared to wildtype (average speed = 0.52±0.01 μm/min; total distance = 341.35±12.86 μm; Fig 4I, J), while net displacement remains unchanged (wildtype=78.50±15.07 μm, vs. *tfap2a;foxd3*=71.48±20.79 μm).

These results reveal that the neural crest regulates cell movements within the optic vesicle, and underscore the power of 4-dimensional individual cell tracking. In all cell populations tracked, we identified differences in cell speed and distance travelled when comparing wildtype to *tfap2a;foxd3* double mutants; additionally, we observed trajectory timing changes in the absence of neural crest indicating delays in rim involution. Most strikingly, however, was the effect of loss of neural crest on the trajectory and positioning of the nasal RPE. While the other cell populations we tracked reach the correct position and do not show differences in net displacement, the cells which comprise the nasal RPE do not reach the correct position in *tfap2a;foxd3* double mutants, as indicated by the change in net displacement. Consistent with neural crest cells migrating in close proximity to the medial side of the optic vesicle (Fig 1) where cells will either contribute to the RPE or undergo rim involution to contribute to the neural retina, these data demonstrate that the neural crest is critical for regulation of RPE cell movements and rim involution during optic cup morphogenesis.

### Basement membrane formation is disrupted around the RPE in *tfap2a;foxd3* double mutants

Having demonstrated a role for neural crest cells in regulating cell movements within the developing optic cup, we sought to determine the underlying molecular mechanism. Although TGF-β signaling had been a tantalizing candidate (Fuhrmann et al., 2000; Grocott et al., 2011), our phospho-Smad3 antibody staining results indicated that TGF-β signaling to the zebrafish optic cup is not disrupted in the absence of neural crest. Having identified defects in cell movements in many regions of the optic cup, we asked how neural crest might regulate these cell behaviors. Prior work in other systems indicated that mesenchymal cells can regulate epithelial morphogenesis via alterations to the extracellular matrix. We therefore asked whether there might be defects in the ECM surrounding the optic cup when neural crest is lost. By using transmission electron microscopy to directly observe ECM, we visualized assembled basement membranes at the basal surfaces of the brain and RPE, as well as the neural retina and lens in 24 hpf control and *tfap2a;foxd3* double mutant embryos (Fig 5). Although neural crest cells migrate in the space between the brain and the optic vesicle, the basement membrane lining the developing forebrain in *tfap2a;foxd3* double mutants appears indistinguishable from control embryos (Fig 5A, B); the basement membranes lining the neural retina (Fig 5E, F) and lens (Fig 5G, H) also appeared normal in the *tfap2a;foxd3* double mutants compared to wildtype controls. To our surprise, we found that only the basement membrane surrounding the RPE was defective in the *tfap2a;foxd3* double mutant (Fig 5D, D’); this basement membrane appears disorganized and discontinuous compared to the same structure in controls (Fig 5C, C’). These results indicate that neural crest cells are required for basement membrane formation around the optic cup, specifically the basement membrane surrounding the RPE. To our knowledge, this is the first indication of neural crest being required to build basement membrane around the developing eye.

**Figure 5.**
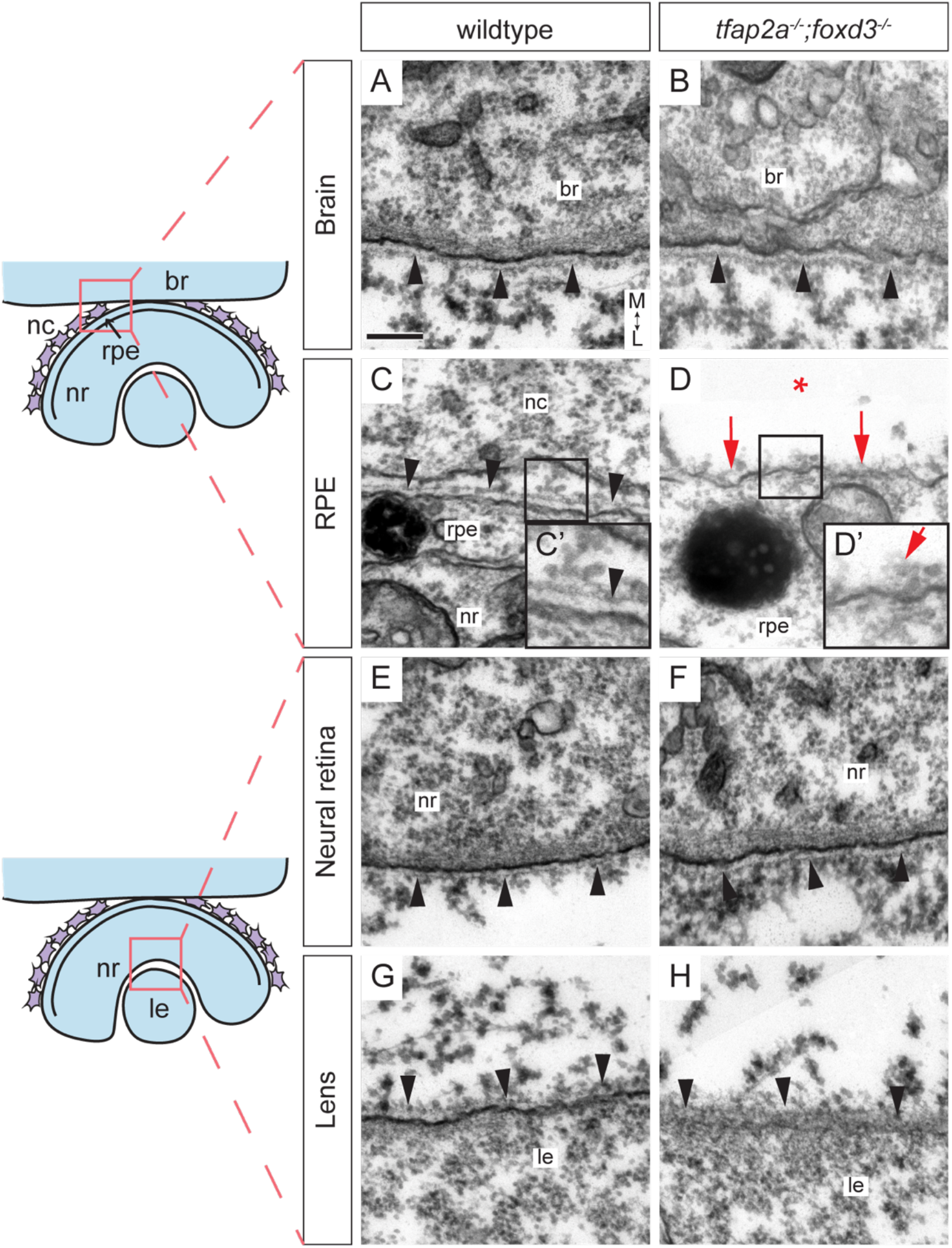
The basement membrane around the RPE is disrupted in *tfap2a;foxd3* double mutants. Transmission electron microscopy was used to visualize the basement membranes around the brain, RPE, neural retina, and lens in 24 hpf control (A, C, E, G) and *tfap2a;foxd3* double mutant (B, D, F, H) embryos, as diagrammed. The basement membrane around the RPE of *tfap2a;foxd3* mutant embryos appears disorganized (D, D’ red arrows) compared to wildtype (C, C’), while all other basement membranes appear normal (black arrowheads). Scale bar, 200 nm. Magnification in (A-H)=10,000x, (C’, D’)=20,000x. br, brain; nc, neural crest cell; nr, neural retina; rpe, retinal pigment epithelium; le, lens. All images are transverse sections, anterior views. M, medial; L, lateral.

### The ECM crosslinking protein Nidogen is produced by neural crest and is lost in the *tfap2a;foxd3* double mutant

To determine how basement membrane formation is disrupted in the *tfap2a;foxd3* double mutant, we next set out to determine what individual extracellular matrix components might be affected. We asked whether localization or expression of the ECM components laminin-1 and fibronectin were altered in the *tfap2a;foxd3* double mutant. Antibody staining for laminin (Fig S3A-D) and fibronectin (Fig S3E-H) revealed no obvious differences between wildtype and double mutant embryos.

In mouse, the ECM protein nidogen has been found to be necessary for lung and kidney epithelial morphogenesis, yet is provided by surrounding mesenchymal cells (Bader et al., 2005; Ekblom et al., 1994; Kadoya et al., 1997; Willem et al., 2002). Nidogen is an extracellular matrix crosslinking protein which is critical for basement membrane assembly in other systems (Bader et al., 2005; Böse et al., 2006; Ekblom et al., 1994; Kadoya et al., 1997), and, along with laminin-1, is one of two ECM proteins required to be included in the medium to support embryonic-stem-cell-derived optic cup morphogenesis *in vitro* (Eiraku et al., 2011). We asked whether nidogen localization might be disrupted around the eye in the *tfap2a;foxd3* double mutant. Antibody staining in wildtype embryos revealed that nidogen protein is detected around cells which exhibit mesenchymal morphology and occupy the same space as neural crest cells migrating between the brain and the optic vesicle at 18 hpf (Fig 6A, A’). Staining is also detectable at the basal surfaces surrounding the RPE, neural retina, and lens at 24 hpf (Fig 6C-C’’), consistent with localization within the basement membrane. In contrast, *tfap2a;foxd3* double mutants that lack neural crest cells also lack nidogen protein between the brain and RPE (Fig 6B, B’, D, D’), yet nidogen staining at the basal surfaces of the lens and at the lens-retina interface is unaffected (Fig 6D’’). These data indicate that nidogen is broadly localized to the ECM in wildtype embryos, but is specifically lost around the RPE when neural crest is lost.

**Figure 6.**
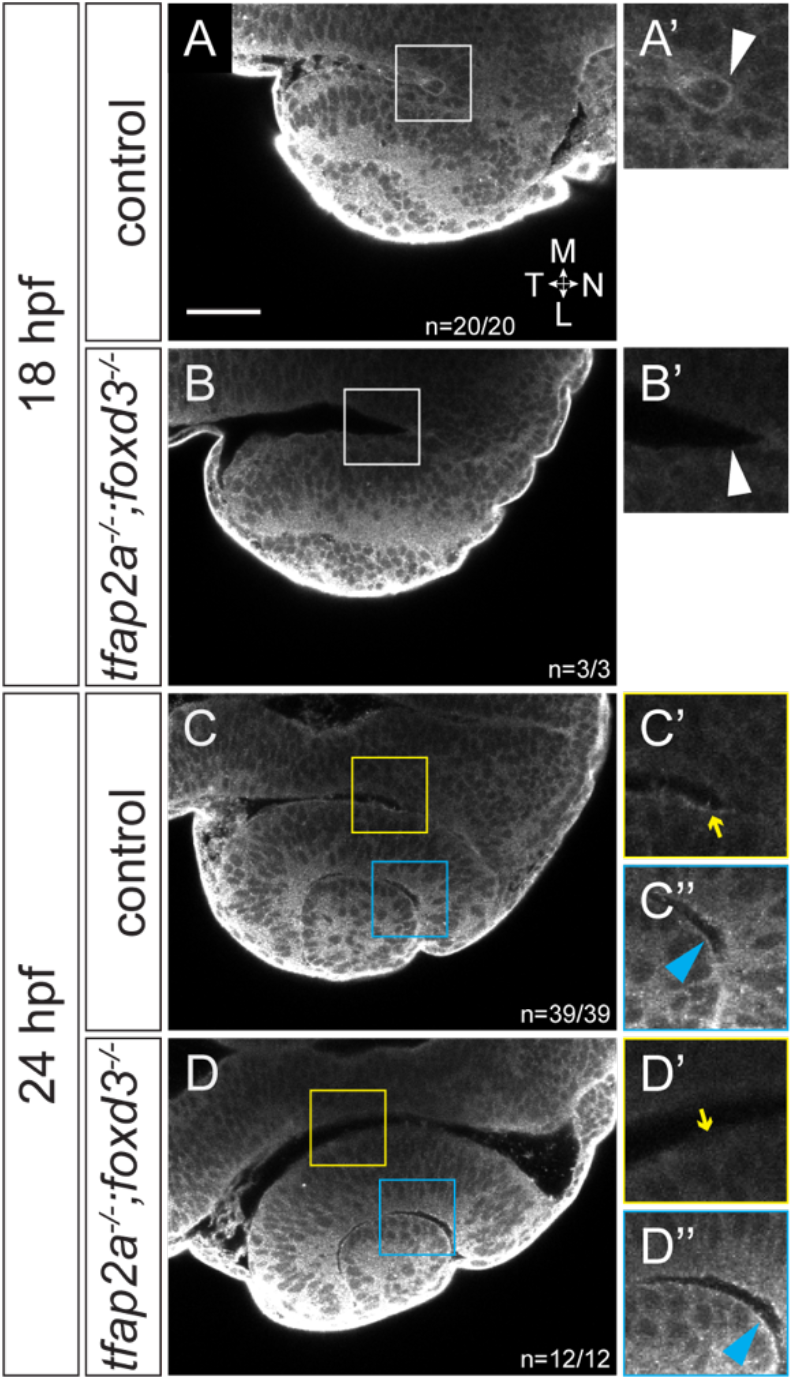
Nidogen protein is absent from the RPE side of the optic cup in *tfap2a;foxd3* double mutants. At 18 hpf, nidogen protein is detected by immunofluorescence in neural crest cells between the brain and developing RPE in the optic stalk furrow (A, magnified in A’). This expression is missing in *tfap2a;foxd3* double mutant embryos at 18 hpf and there are no cells visible in the same space (B, magnified in B’). At 24 hpf in control embryos, a nidogen ECM surface is detectable on along both the RPE (yellow box in C, yellow arrow in C’) and at the lens-retina interface (C, C’’, blue arrowhead). In 24 hpf *tfap2a;foxd3* double mutants, nidogen is not detectable along the RPE (D’, yellow arrow), but is still present in the ECMs at the lens-retina interface (D’’, blue arrowhead). Dorsal view, single confocal sections. Scale bar, 50 μm.

In mouse, nidogen expression has been observed in the POM and invaginating lens (Dong and Chung, 1991), and zebrafish *in situ* hybridizations suggest that both *nidogen 1b* and *nidogen 2a* are expressed in the cranial neural crest (Kudoh et al., 2001; Thisse and Thisse, 2004; Zhu et al., 2017). Despite these observations, the role of nidogen during optic cup morphogenesis has not been investigated *in vivo*. Because the specific periocular cell populations expressing nidogens had not been completely defined, we sought to determine with cellular resolution the expression of the four zebrafish *nidogen* genes in and around the developing eye. Using *Tg(sox10:GFP)^ba4^* transgenics, we found that both *nid1b* and *nid2a* are expressed in *sox10:GFP*- positive neural crest cells migrating around the developing optic cup (Fig 7). We also observed both *nid1b* and *nid2a* expressed in the overlying ectoderm and the developing lens at 18 and 24 hpf, while both are notably absent from the neural retina and RPE. At these same times, *nid1a* is expressed solely in the developing somites (Fig S4A, E, I), while *nid2b* is detected diffusely throughout the head (Fig S4D, H, L).

**Figure 7.**
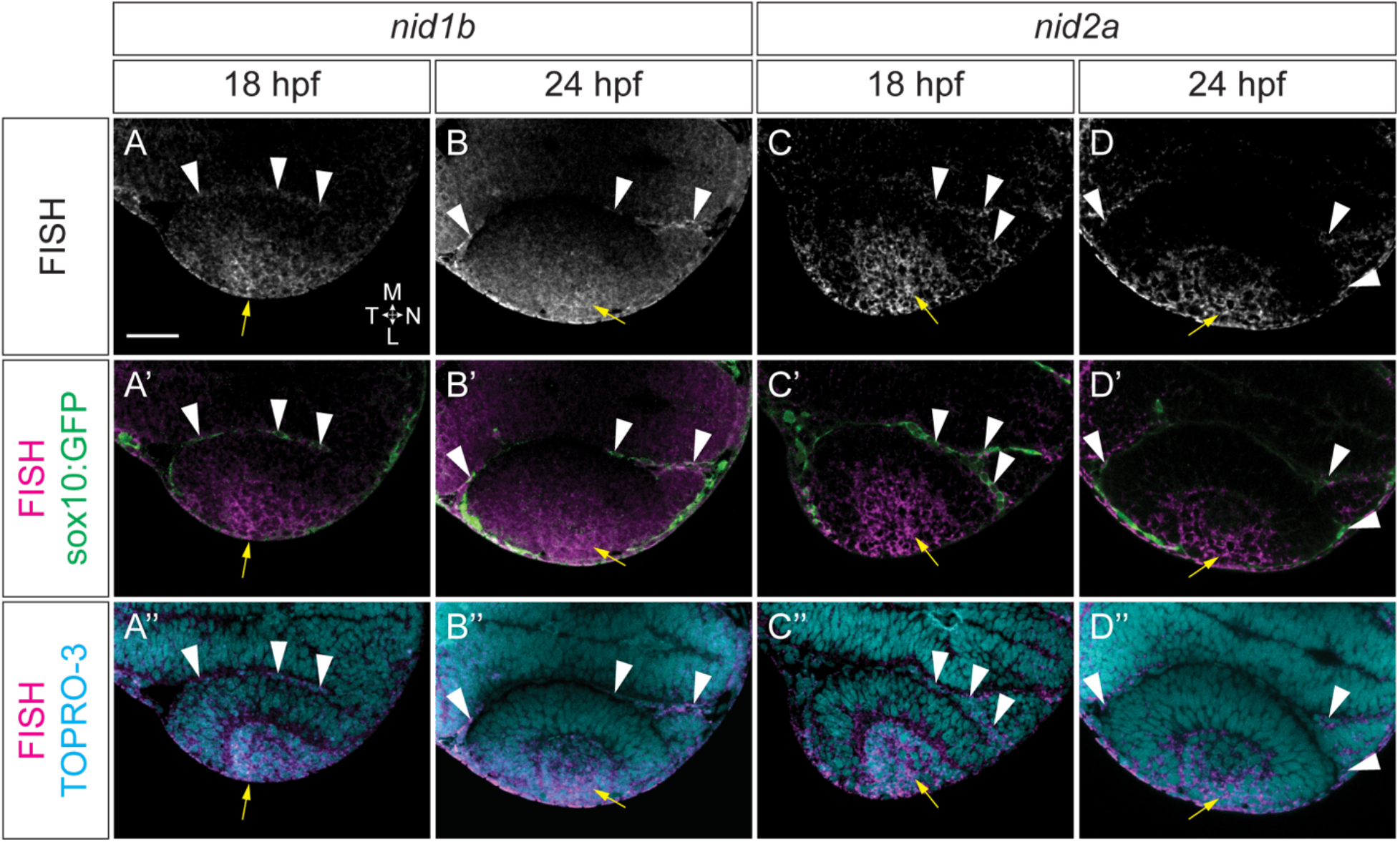
Nidogen 1b and 2a are expressed in the neural crest and developing lens. *In situs* were performed in *Tg(sox10:GFP)* embryos at 18 (A-A’’, C-C’’) and 24 hpf (B-B’’, D-D’’). (A-D) Fluorescence *in situ* hybridization with probes against *nid1b* (A,B) and *nid2a* (C,D). (A’-D’) FISH merged with sox10:GFP expression (green) to visualize colocalization between FISH and GFP^+^ neural crest cells (white arrowheads). GFP signal was amplified after hybridization using an anti-GFP antibody. (A’’-D’’) FISH merged with nuclei counterstained with TO-PRO-3. Yellow arrows denote lens expression in each case. Dorsal view, single confocal sections. Scale bar, 50 μm.

Taken together, these data suggest that neural crest cells produce the nidogen proteins found in the basal surface of the developing RPE, while the ectoderm and lens placode produce nidogen found surrounding the lens and basal surface of the neural retina.

### Dominant-interfering nidogen disrupts optic cup morphogenesis

The basement membrane loss we observed around the RPE in the *tfap2a;foxd3* double mutant was reminiscent of the basement membrane loss observed when nidogen has been disrupted in mouse models of organogenesis, either through loss of function mutations or through disruptions to laminin-nidogen complex formation (Bader et al., 2005; Ekblom et al., 1994; Kadoya et al., 1997; Willem et al., 2002). This observation, coupled with the finding that *nid1b* and *nid2a* are expressed by neural crest cells during optic cup morphogenesis, suggested a potential functional role for nidogen in optic cup morphogenesis. Previous studies have reported genetic compensation when some but not all nidogens were disrupted (Bader et al., 2005; Böse et al., 2006; Zhu et al., 2017); since there are two nidogens expressed by neural crest and a third expressed diffusely throughout the head, we elected to use a dominant-interference strategy to disrupt nidogen function. We took advantage of a well characterized truncated form of nidogen, Nd-III, which lacks the G1 and G2 domains required for binding to collagen IV and heparin sulfate proteoglycan, but retains the laminin binding domain (Fox et al., 1991; Reinhardt et al., 1993). As shown through characterization *in vitro*, Nd-III acts in a dominant-inhibitory fashion by blocking bridging between laminin and other extracellular matrix components, and interfering with full length nidogen binding to laminin (Pujuguet et al., 2000). We generated transgenic zebrafish which expressed lyn-mCherry as a reporter, as well as a viral 2A peptide upstream of either full-length zebrafish *nid1a* (Fig 8A, WT-Nid1a), or a truncated form of *nid1a* based on the mouse Nd-III fragment (Fig 8A, DI-Nid1a). By using heat-shock inducible transgenes, we were able to control the timing of expression of WT-Nid1a and DI-Nid1a, and by driving lyn-mCherry from the same transcript, we controlled for efficiency of heat shock by visualizing all cells which expressed the transgene (see Materials and Methods for full transgenic nomenclature).

**Figure 8.**
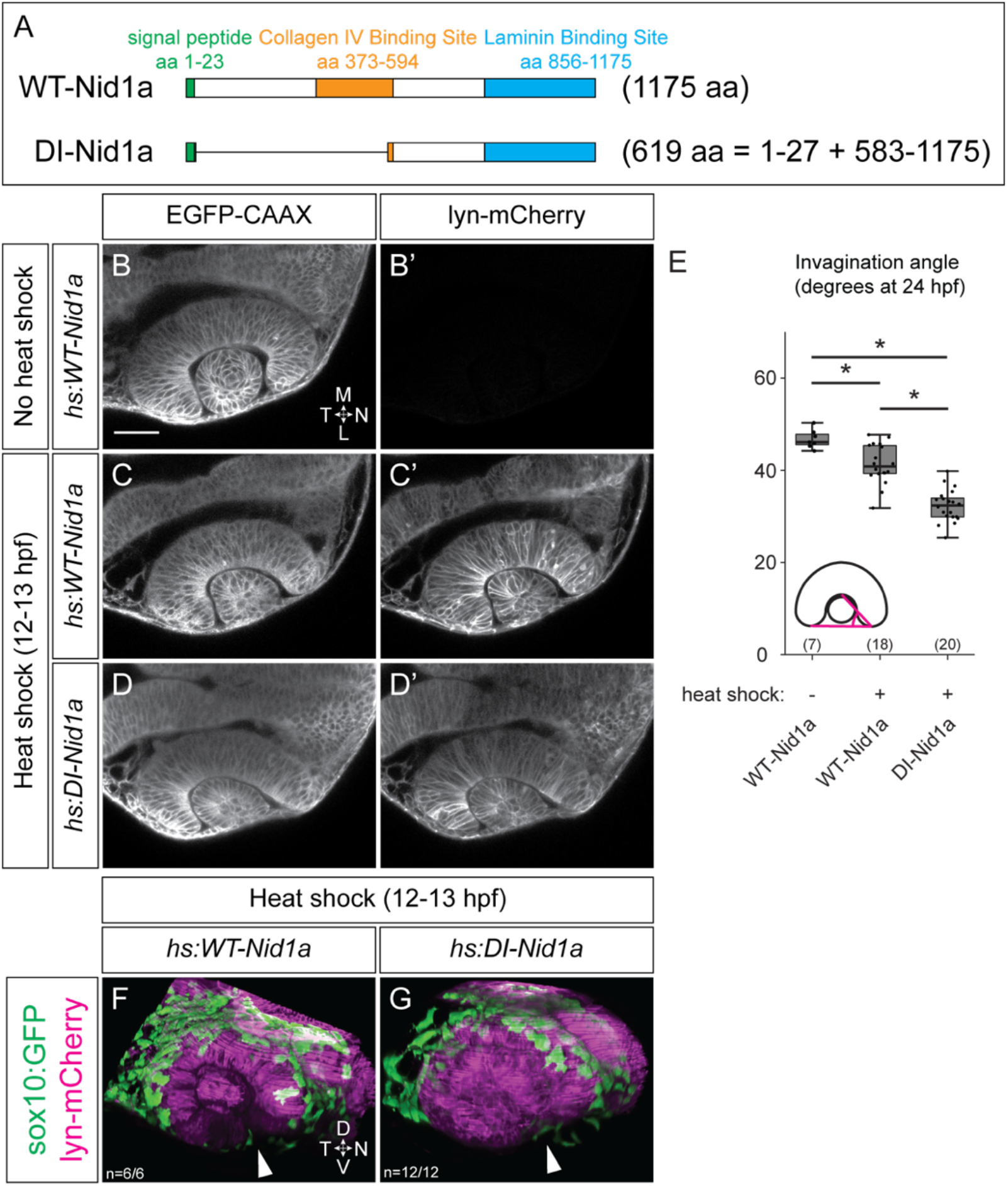
Dominant-interfering nidogen disrupts optic cup morphogenesis. (A) Schematics of full length (WT) and dominant-interfering (DI) Nidogen 1a. (B-D’) Dorsal view, single confocal sections. *Tg(bactin2:EGFP-CAAX)* females were crossed to either *hs:WT-Nid1a* (B-C’) or *hs:DI-Nid1a* (D, D’) transgenic males. Control embryos (B, B’) were not heat shocked, show no lyn-mCherry expression; experimental embryos were heat shocked 12-13 hpf (C-D’). (E) Quantification of invagination angles, measured as shown in inset diagram. ^*^P<0.005 using Welch’s t-test. (*WT-Nid1a* no heat shock vs. heat shock, P=0.001; *WT-Nid1a* heat shock vs. *DI-Nid1a* heat shock, P=4.34×10^-10^; *WT-Nid1a* no heat shock vs. *DI-Nid1a* heat shock, P=7.76×10^-11^; *DI-Nid1a* no heat shock vs. heat shock (not shown), P=6.46× 10^-14^.) Results are from 3 experiments; n (embryos) shown in base of graph. (F, G) Lateral view, 3D renderings at 24 hpf. *sox10:GFP* transgenic females were crossed to either *hs:WT-Nid1a* (F) or *hs:DI-Nid1a* (G) transgenic males, embryos heat shocked 12-13 hpf. GFP-positive neural crest cells migrate around the optic cup and into the optic fissure (white arrowheads) in both conditions. Scale bar, 50 μm. M, medial; L, lateral; N, nasal; T, temporal.

We previously observed neural crest cells in contact with the optic vesicle as early as 12.5 hpf (Fig 1), therefore we performed heat shock from 12-13 hpf to induce WT-Nid1a or DI-Nid1a expression at the onset of neural crest migration around the optic vesicle. We crossed zebrafish containing *hs:WT-Nid1a* or *hs:DI-Nid1a* transgenes with *Tg(bactin2:EGFP-CAAX)^z200^* transgenics; this enabled us to image the optic cups of *hs:WT-Nid1a* transgene-positive embryos which had not been heat shocked (Fig 8B, B’) as a control. We found that ubiquitous overexpression of WT-Nid1a slightly, but significantly, impaired invagination (42.3±1.7°) compared to control embryos (46.8±1.4°; Fig 8C, C’, E), and subtly altered lens morphogenesis such that the resulting lens is not completely spherical. However, overexpression of DI-Nid1a significantly impaired invagination (32.3±1.5°), resulting in striking phenotypes including a severely flattened neural retina and lens (Fig 8 D, D’, E). To determine whether these phenotypes were due to effects on the optic cup itself or a secondary consequence of disruption to neural crest migration, we generated double transgenics with *sox10:GFP* and either *hs:WT-Nid1a* or *hs:DI-Nid1a* to visualize neural crest cells in the presence of WT-Nid1a or DI-Nid1a. We find that ubiquitous, heat shock-induced overexpression of WT-Nid1a or DI-Nid1a does not affect neural crest migration (Fig 8F, G). We conclude, therefore, that DI-Nid1a strongly impairs optic cup morphogenesis through direct effects on the optic vesicle.

To determine the effects of disrupting nidogen specifically expressed by the neural crest, we attempted to achieve neural crest-specific expression of either WT-Nid1a or DI-Nid1a by utilizing the Gal4/UAS system. We generated *UAS:WT-Nid1a* and *UAS:DI-Nid1a* transgenic lines and crossed each to *Tg(sox10:Gal4-VP16)^el159^* transgenic zebrafish (Das and Crump, 2012). Unlike ubiquitous, heat-shock induced overexpression, neural crest-specific overexpression of WT-Nid1a did not have an effect on optic cup morphogenesis (data not shown). While neural crest-specific expression of DI-Nid1a (*sox10:Gal4-VP16;UAS:DI-Nid1a* double transgenic) led to obvious optic cup invagination defects, these embryos had very few or no neural crest cells in the vicinity of the eye (data not shown), either due to effects on neural crest survival or migration. Thus we were unable to distinguish the effects of DI-Nid1a directly on the eye from effects which arise due to loss of neural crest. We hypothesize that while a short burst of DI-Nid1a expression (such as when induced from 12-13 hpf via heat shock) does not impair neural crest formation or migration (Fig 8G), sustained expression of DI-Nid1a (as driven by the *sox10* promoter) impairs neural crest migration and/or survival, and thus we were unable to interpret the tissue-specific effects of DI-Nid1a expression from the neural crest.

### Nidogen can partially rescue *tfap2a;foxd3* double mutant optic cup phenotypes

Nidogen expression in the neural crest, coupled with the phenotypes we observed when we ubiquitously overexpressed DI-Nid1a, suggested that disruptions to nidogen’s matrix bridging function might be the underlying cause of the morphogenesis defects we observed in mutants where neural crest cells are lost.

We sought to determine whether the optic cup morphogenesis defects we observed in the *tfap2a;foxd3* mutants might indeed be due to a lack of nidogen deposited by neural crest cells. Specifically, we wanted to determine whether expression of WT-Nid1a could rescue optic cup morphogenesis in *tfap2a;foxd3* double mutants. To test this, we ubiquitously overexpressed WT-Nid1a in *tfap2a;foxd3* double mutants where there would be no neural crest cells present to produce nidogen. We generated *tfap2a;foxd3* double mutants that contained both the *EGFP-CAAX* transgene, as well as the *hs:WT-Nid1a* transgene. In these embryos, we could visualize the optic cups of both unperturbed embryos, as well as those which were heat-shocked from 12-13 hpf. In control, non-heat shocked embryos (Fig 9A-B’), we observed retinal invagination defects in *tfap2a;foxd3* double mutants compared to their wildtype siblings (Fig 9E), consistent with our previous observations (Fig 2G). To our surprise, wildtype and *tfap2a;foxd3* mutant embryos that were heat shocked from 12-13 hpf looked phenotypically similar. The lenses in either genotype take on a similar, slightly ovoid appearance, and the nasal retina more fully enwraps the lens in *tfap2a;foxd3* mutants which overexpress WT-Nid1a, suggesting improved invagination over *tfap2a;foxd3* mutant eyes (Fig 9C-D’). When quantified, we indeed observe a significant improvement in the degree of invagination between *tfap2a;foxd3* double mutant eyes (36.8±1.6°) and *tfap2a;foxd3* double mutants which ubiquitiously overexpress WT-Nid1a (39.9±1.8°; Fig 9E). Additionally, we did not detect a difference between heat shocked wildtype and *tfap2a;foxd3* embryos when quantifying invagination (39.6±0.7° vs 39.9±1.8°, respectively; Fig 9E). From this experiment, we conclude that supplying nidogen can partially bypass the effects of loss of neural crest on optic cup morphogenesis, and that ectopic expression of nidogen from the optic cup and surrounding tissues can partially rescue the invagination defects caused by loss of nidogen producing cells.

**Figure 9.**
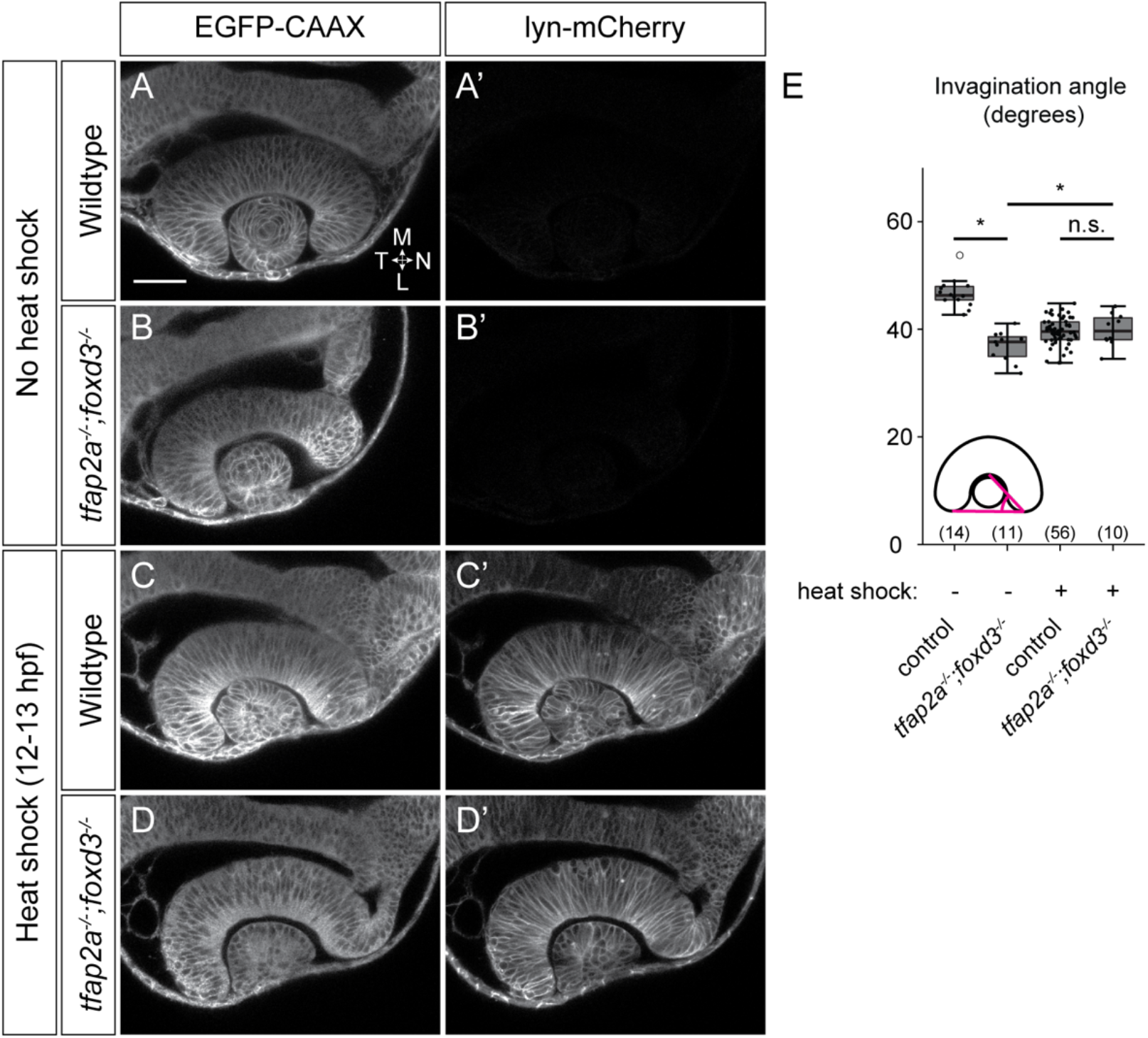
Overexpression of Nid1a partially rescues optic cup morphogenesis in *tfap2a;foxd3* double mutants. Dorsal view, single confocal sections from 24 hpf *Tg(hs:WT-Nid1a)*-positive embryos.(A-B’) EGFP-CAAX and lyn-mCherry channels from wildtype (A, A’) and *tfap2a;foxd3* double mutant (B, B’) embryos which were not subjected to a heat shock. (C-D’) EGFP-CAAX and lyn-mCherry channels from wildtype (C, C’) and *tfap2a;foxd3* double mutant (D, D’) embryos which were heat shocked from 12-13 hpf. (E) Quantification of invagination angles, measured as shown in the inset diagram. ^*^P<0.05 using Welch’s t-test; n.s., not significant. (Wildtype (no heat shock) vs. *tfap2a;foxd3* double mutant (no heat shock), P=1.42× 10^-8^; *tfap2a;foxd3* double mutant no heat shock vs. heat shock, P=0.03; wildtype (heat shock) vs. tfap2a;foxd3 (heat shock), P=0.76.) Results are from 3 experiments; n (embryos) shown in the base of the graph. Scale bar, 50 μm. M, medial; L, lateral; N, nasal; T, temporal.

## Discussion

### Interactions with neural crest cells regulate optic cup morphogenesis

Crosstalk between developing tissues is critical for their proper morphogenesis and subsequent function. This theme is seen throughout organogenesis, with epithelial-mesenchymal interactions being of particular importance. Developing epithelia frequently require surrounding mesenchyme in order to acquire their mature, functional structures: the lung, kidney, salivary gland, and tooth placode are but a few examples where these interactions are critical (Bader et al., 2005; Ekblom et al., 1994; Kadoya et al., 1997; Thesleff, 2003). In the eye, a complex mesenchyme is known to surround the developing optic cup, and mesenchymal cells contribute to later-developing optic tissues such as the hyaloid vasculature and structures within the anterior segment (Gage et al., 2005; James et al., 2016; Soules and Link, 2005; Sowden, 2007). Prior work has established that disruptions to the POM have profound effects on the developing optic cup, such as morphogenetic defects that stretch from the neural retina into the optic stalk and optic fissure (Bassett et al., 2010; Lupo et al., 2011). However, many of these analyses leave many questions unresolved, such as when or where defects arise within the optic cup. The POM is a heterogeneous cell population comprised of both neural crest and mesodermally-derived mesenchymal cells (Gage et al., 2005; Johnston et al., 1979), and specific molecular contributions from either tissue during early optic cup development are poorly understood. A recently published study using elegant ectopic optic vesicle transplants suggested that the POM was dispensable for early optic cup morphogenesis and only required for later stages of optic cup development, specifically the fusion of the optic fissure (Gestri et al., 2018). Here, using the zebrafish *tfap2a;foxd3* double mutant which displays a near complete loss of neural crest cells (Arduini et al., 2009; Wang et al., 2011), we demonstrate that the neural crest subpopulation of the POM regulates morphogenesis of the optic cup. Using live imaging, we visualize neural crest cells and find that they migrate around the optic vesicle throughout much of optic cup morphogenesis. Despite relatively mild gross morphological defects, 4-dimensional cell tracking reveals that neural crest cells are required for cell movements within optic vesicle. Specifically, proper RPE cell movements and rim involution depend on the presence of neural crest cells; in the absence of neural crest, cells which contribute to the nasal retina and RPE move significantly farther and faster than in the presence of neural crest. Most notable, however, is the dramatic loss of basement membrane surrounding the RPE when neural crest is lost, which is likely to underlie these cell movement defects (Figure 10).

**Figure 10.**
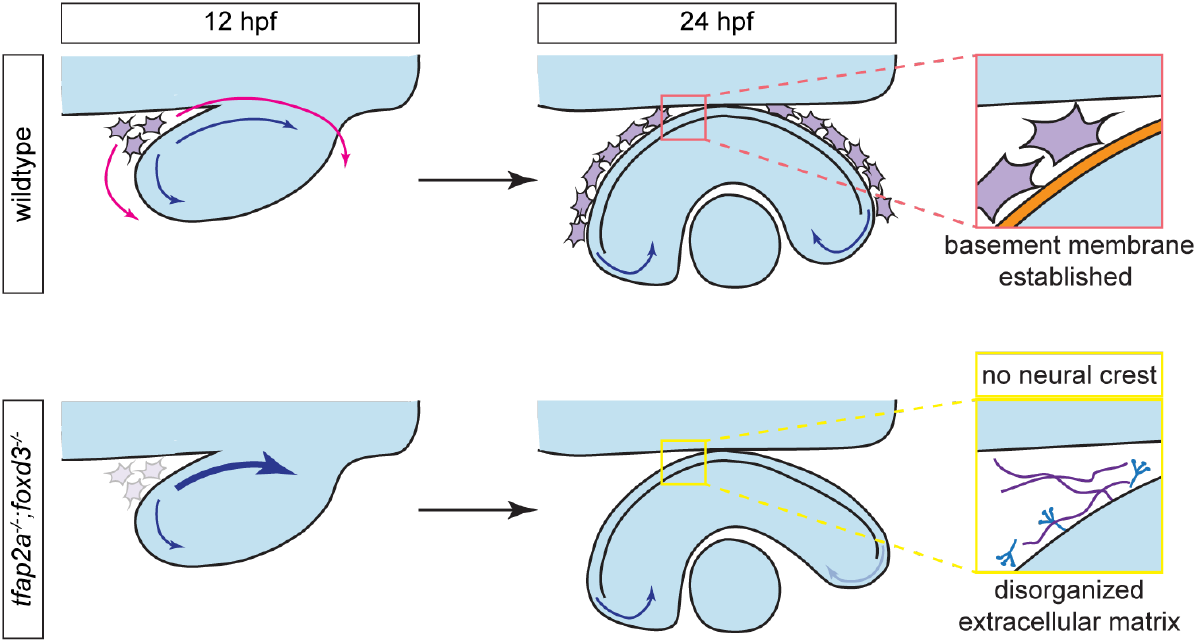
Model of neural crest function during optic cup morphogenesis. Optic cup morphogenesis in a wildtype embryo (top row) and a *tfap2a;foxd3* double mutant (bottom row). In wildtype embryos, neural crest cells (purple) begin to migrate around the optic vesicle (magenta arrows) starting around 12 hpf and enwrap the retinal pigment epithelium by 24 hpf. During this process neural crest cells express and deposit nidogen, which establishes the basement membrane (orange) around the RPE and restricts cell movements within the prospective eye (blue arrows), in turn enabling proper rim movement and optic cup morphogenesis. In *tfap2a;foxd3* double mutants which lack most neural crest cells, the ECM surrounding the RPE is disorganized and does not form a coherent basement membrane. Cell movements within the optic vesicle are unrestricted (heavy blue arrow), leading to the abnormal position of the anterior RPE and disrupted rim movement of cells which would normally migrate into the neural retina.

The POM has long been observed in close proximity with the developing optic cup, but its specific functions have remained elusive. One possible mechanism mesenchymal cells may use to drive optic cup morphogenesis is by modulating the signaling received by the optic vesicle during its development. Zebrafish *zic2a;2b* double mutants display disruptions to the periocular neural crest, in addition to molecular and morphological hallmarks of expanded Hedgehog signaling within the optic cup (Sedykh et al., 2017); in *tfap2a;foxd3* double mutants however, we do not observe Pax2a (a Hh target gene) expression expanded in a manner consistent with disrupted Hh signaling. In chick, periocular mesenchyme is required for RPE development in a seemingly TGF-β dependent fashion (Fuhrmann et al., 2000). Using pSmad3 as a readout, we did not observe a difference in TGF-β signaling in the optic cup in *tfap2a;foxd3* double mutants; additionally, neural crest cells are not required for the development of RPE in zebrafish, as we observed pigment granule formation by electron microscopy at 24 hpf, and obvious pigmentation of *tfap2a;foxd3* mutant eyes at 52 hpf. It is possible that that the mesodermal mesenchyme is sufficient for TGF-β signaling to the optic cup and RPE induction, or that RPE development could be regulated through different mechanisms in the zebrafish versus the chick. In the future, it will be interesting to dissect the specific roles of the mesodermal mesenchyme in optic cup morphogenesis.

An alternative explanation for the phenotypes we observe in the *tfap2a;foxd3* double mutants is that loss of either gene could affect other surrounding tissues in close proximity to the eye, in addition to their well-characterized effects on the neural crest. *tfap2a* in particular is expressed in the overlying ectoderm during optic cup morphogenesis (Bassett et al., 2010; Knight et al., 2005). As such, its loss could affect some aspect of the optic cup morphogenesis through a neural crest-independent effect. However, both *tfap2a* and *foxd3* single mutants display optic cup morphogenesis defects that are less severe than those we observe in the *tfap2a;foxd3* double mutant. In addition, the lens placode is the sole optic tissue where *tfap2a* is expressed during optic cup morphogenesis stages, yet the lens is morphologically normal in each of these mutants. Further, we observe disruptions to basement membrane formation along the surface of the RPE which is in direct contact with neural crest cells, but normal basement membrane formation at the basal surface of the retina and lens. These observations suggest that the optic cup morphogenesis defects we observe in the *tfap2a;foxd3* double mutants are primarily due to loss of neural crest cells rather than effects on other surrounding tissues.

### Neural crest cells promote morphogenesis through ECM assembly

Several studies have implicated the ECM surrounding the optic vesicle as a key player in driving optic cup morphogenesis. Fibronectin appears to be critical for formation of the lens (Hayes et al., 2012; Huang et al., 2011), and in retinal organoid culture, the minimal mixture of components needed to elicit optic cup morphogenesis are nodal (a TGF-β ligand) and the ECM components laminin-1 and nidogen (Eiraku et al., 2011). Laminin is required for proper cell adhesion of the optic vesicle to the ECM, and disruptions to these adhesions impairs establishment of apicobasal polarity and subsequent morphogenesis (Bryan et al., 2016; Ivanovitch et al., 2013; Nicolás-Pérez et al., 2016; Sidhaye and Norden, 2017). Taken together, these data indicate that the ECM is a critical regulator in formation of the optic cup, and the individual components which comprise the ECM each may regulate specific, non-overlapping aspects of this morphogenetic process.

Further, the dynamics of ECM component production, deposition, and interaction may play a key role in regulating cell movements. Using electron microscopy, we observe a spatially specific, dramatic disruption of basement membrane formation when neural crest is lost. This has allowed us the opportunity to determine the effect of disrupting ECM assembly in a limited domain, without loss of key the basement membrane component laminin. The gross morphology of the optic cup is mildly but reproducibly abnormal, although cell movements in contact with that particular ECM domain are clearly disrupted. We speculate that the intact basement membrane in other parts of the eye is sufficient to support other cell movements underlying optic cup morphogenesis. That being said, comparison of this phenotype with other mutants which disrupt ECM components reveals some notable findings: for example, loss of laminin leads to significant cell death in the prospective RPE domain (Bryan et al., 2016); impairment of basement membrane formation through genetic loss of neural crest cells does not. Further examination of these phenotypes will allow us to identify functions of ECM components that are specifically dependent upon basement membrane assembly. It is also interesting to consider that cells within the developing RPE may rely on the mechanical properties of the surrounding basement membrane in order to migrate and develop correctly. The Hippo signaling pathway is a major cellular mechanotransducer which can function in response to ECM stiffness (Chakraborty et al., 2017), and RPE development is dependent on Yap and Taz, the transcriptional coactivators of the Hippo pathway (Miesfeld et al., 2015). Although RPE development appears normal without assembled basement membrane, it is a tantalizing possibility that Hippo signaling could serve as a molecular link between the basement membrane and optic vesicle cell behaviors.

Although the role of nidogen in other organ systems has been previously investigated, its roles in optic cup morphogenesis has remained elusive until now. While nidogens are ubiquitous components of basement membranes, the mesenchymal cells in the embryonic lung and kidney are those tissues’ sole source of nidogens (Ekblom et al., 1994; Kadoya et al., 1997; Senior et al., 1996). Disruptions to nidogen function, either through blocking antibodies or loss-of-function mutations, impedes epithelial morphogenesis of the lung, kidney and developing limb (Bader et al., 2005; Böse et al., 2006; Ekblom et al., 1994; Kadoya et al., 1997). Here, we demonstrate that neural crest cells produce the nidogen which is deposited along the basal surface of the RPE, and that neural crest cells are required for proper assembly of the basement membrane along this surface. Impairing nidogen function, either through loss of neural crest cells or through expression of a dominant-interfering form of Nid1a, leads to defects in optic cup morphogenesis. Intriguingly, ubiquitous overexpression of full-length Nid1a also causes slight defects in optic cup invagination, suggesting that an optimal level of nidogen is required for proper ECM function. Consistent with this, loss of the nidogen-binding site in the laminin *γ*1 chain results in higher penetrance of renal defects than when both nidogen 1 and 2 are lost (Bader et al., 2005; Willem et al., 2002); this was predicted to be due to an increase of free nidogen incapable of binding to laminin, which could sequester other components of the ECM and interfere with other matrix interactions (Bader et al., 2005).

How the addition of nidogen to the ECM surrounding the optic cup promotes the morphogenetic movements required for optic cup morphogenesis is unclear. While both laminin and fibronectin were still detectable around the RPE in the absence of neural crest cells, we did not detect a normal basement membrane at this site. This finding is consistent with mouse nidogen mutants which show similar basement membrane defects around the developing kidney, lung, and heart, despite retention of other ECM components (Bader et al., 2005). Proper attachment to the ECM is required for many aspects of optic cup morphogenesis including RPE cell movements and rim involution (Bryan et al., 2016; Hayes et al., 2012; Martinez-Morales et al., 2009; Sidhaye and Norden, 2017), two processes which we show to be disrupted in the absence of neural crest. We propose that through deposition of nidogen into the ECM, neural crest cells generate a restrictive environment which enables the correct movements of RPE cells and migration of the cells which move around the rim of the optic vesicle and into the developing neural retina; the basement membrane surrounding the RPE serves as a handbrake which slows optic vesicle cell movements and ensures their migration to the correct place at the correct time during optic cup morphogenesis. Disruptions to nidogen function, either through loss of neural crest cells, expression of a dominant-interfering form of nidogen, or overexpression of wildtype nidogen, lead to defects in optic cup morphogenesis. It will be interesting to investigate whether cells in the developing optic cup can still properly adhere to the remaining ECM components in the absence of neural crest cells. Further, growth factors and signaling molecules such as FGFs and BMPs are regulated through deposition into the ECM, and correct assembly of the basement membrane may be required to properly regulate these signaling pathways during optic cup morphogenesis; these will be interesting signaling pathways to investigate in the context of disrupted basement membrane assembly.

## Materials and methods

### Zebrafish lines

Embryos from the following mutant and transgenic lines were raised at 28.5°C and staged according to hours post fertilization and morphology (Kimmel et al., 1995).

#### Mutant alleles

*tfap2a^ts213^;foxd3^zdf10^* (Arduini et al., 2009). The *alyron^z24^* allele contains a C to A transversion in the coding sequence of the *paf1* gene, resulting in a premature stop mutation at tyrosine 281 (Y281^*^) (Mick Jurynec and David Grunwald, personal communication).

#### Transgenic alleles

*Tg(sox10:memRFP) ^vu234^* (Kirby et al., 2006), *Tg(sox10:GFP)^ba4^* (Dutton et al., 2008), *Tg(bactin2:EGFP-CAAX)^z200^*, *Tg(hsp70:lyn-mCherry-2A-WT-Nid1a)^z202^, Tg(hsp70:lyn-mCherry-2A-DI-Nid1a)^z203^.*

### Construction of Nid1a transgenic lines

*Tg(hsp70:lyn-mCherry-2A-WT-Nid1a)*, a.k.a. *hs:WT-Nid1a* and *Tg(hsp70:lyn-mCherry-2A-DI-Nid1a),* a.k.a. *hs:DI-Nid1a* were generated using Gateway (Invitrogen, Carlsbad, CA) recombination. IMAGE Clone ID 8000296 (GE Dharmacon, Chicago, IL) was used as the template to PCR amplify cDNAs encoding wildtype and dominant-interfering Nid1a, these were ligated into pCS2FA prior to Gateway cloning. PCR primers were used to introduce the PTV-2A peptide (Provost et al., 2007) on the 5’ end, and the SV40 late poly-adenylation signal on the 3’ end of the zebrafish *nid1a* cDNA. Gateway 3’ entry clones were generated via BP recombination and subsequently LR recombined into the pDEST-Tol2-CG2 destination vector which contains an *myl7:EGFP* expression cassette as a transgenesis marker (Kwan et al., 2007). 25 pg plasmid DNA was microinjected along with 50 pg mRNA encoding Tol2 transposase into single-cell wildtype embryos and screened for *myl7:EGFP* expression. Fluorescent embryos were raised to adulthood and outcrossed to generate stable transgenic lines.

### Heat shocks

Embryos were transferred from a 28.5°C incubator and immediately overlaid with fresh, preheated 39°C E3. Embryos were incubated at 39°C for one hour on an Echotherm heating plate (Torrey Pines Scientific, Carlsbad, CA). Embryos were then transferred back to a 28.5°C incubator and grown to the indicated stage.

### Allele identification/genotyping

All mutant alleles were PCR genotyped using either CAPS or dCAPS techniques (Neff et al., 1998). *tfap2a^ts213^*: Forward (5’-CGCTCAGGTCTTATAAATAGGCTACTAATAATGTTAC-3’), Reverse (5’-CTGAGAGGTGGCTATTTCCCGTTAAGATTCG-3’), mutant allele is cut with BlpI.

*foxd3^zdf10^*: dCAPS Forward (5’-CGACTGCTTCGTCAAGATCCCACGGGAACCGGGCAACCCGGGCAAAGGCAACTACT GGACCCTCGACCCCCAGTCGGAAAATAT-3’), Reverse (5’-CAGGGGGAATGTACGGGTACTGC-3’), mutant allele is cut with SspI. *paf1^z24^*: Forward (5’-GTTCAGAGGTATGATGGATGAGG-3’), Reverse (5’-GTATGCAGCTTTATGAAAACACTC-3’), wildtype band is cut with NspI.

### RNA synthesis and injections

Capped mRNAs were synthesized using linearized pCS2 templates (pCS2-EGFP-CAAX, pCS2FA-H2A.F/Z-mCherry), the mMessage mMachine SP6 kit (AM1340, Invitrogen), purified using the Qiagen RNeasy Mini Kit (Qiagen, Hilden, Germany) and ethanol precipitated. 150-250 pg of each mRNA were microinjected into the cell of one-cell stage embryos. EGFP-CAAX mRNA was injected to visualize cell membranes, H2A.F/Z-mCherry mRNA was injected to visualize nuclei.

### Antibody staining

Embryos were fixed at the indicated stage in 4% paraformaldehyde, permeabilized in PBST (PBS+0.5% Triton X-100), and blocked in PBST + 2% bovine serum albumin. Antibodies and concentrations are as follows: anti-Pax2a (GTX128127, Genetex, Irvine, CA), 1:200; anti-pSmad3 (ab52903, Abcam, Cambridge, MA), 1:200; anti-Nidogen/Entactin (ab14511, Abcam), 1:100; anti-Laminin 1 (L9393, Sigma, St. Louis, MO), 1:100; anti-Fibronectin (F3648, Sigma), 1:100; anti-GFP (A10262, Invitrogen), 1:200. Secondary antibodies used were Alexa Fluor 488 goat anti-mouse (A-11001, Invitrogen), Alexa Fluor 488 goat anti-rabbit (A-11008, Invitrogen), Alexa Fluor 488 goat anti-chicken (A-11039, Invitrogen) and incubated at 1:200. Nuclei were detected by incubation with 1 μM TOPRO-3 iodide (T3605, Invitrogen). Embryos were cleared through a series of 30%/50%/70% glycerol (in PBS) prior to imaging.

### In situ hybridization

Embryos were fixed at the indicated stage in 4% paraformaldehyde overnight at 4°C and dehydrated in 100% methanol. Color *in situ* hybridizations were performed similar to (Thisse and Thisse, 2008). Fluorescent *in situ* hybridizations were carried out as described previously (Lauter et al., 2014; Leerberg et al., 2017). Anti-GFP labeling and detection was performed after *in situ* hybridization and tyramide signal amplification.

In situ probes were synthesized from linearized pBluescript II SK+ templates (pBSII-Nid1a, pBSII-Nid1b, pBSII-Nid2a, pBSII-Nid2b) using T3 or T7 polymerases and DIG labeling mix (11277073910, Roche, Basel, Switzerland). All four probe sequences were synthesized (IDT gBlocks, IDT, San Jose, CA) and ligated into pBluescript II SK+.

### Light Microscopy

For timelapse imaging, 12 hpf embryos were embedded in 1.6% low-melt agarose (in E3) in DeltaT dishes (Bioptechs, #0420041500 C), E3 was overlaid and the dish covered to prevent evaporation. For antibody stained or fluorescent in situ hybridization imaging, embryos were embedded in 1% low-melt agarose (in PBS) in Pelco glass-bottom dishes (#14027, Ted Pella, Redding, CA), PBS was overlaid to prevent evaporation.

Confocal images were acquired using a Zeiss LSM710 laser scanning confocal microscope. For timelapse imaging, datasets were acquired using the following parameters: 63 z-sections, 2.10 μm z-step, 40x water-immersion objective (1.2 NA). Time between z-stacks was 3 minutes 30 seconds (Movies S1, S2), 2 minutes 45 seconds (Movies S4, S6, S8), and 2 minutes 30 seconds (Movies S5, S7, S9). For all timelapse and antibody imaging, datasets were acquired without knowledge of embryo genotype. Embryos were de-embedded and genotyped after imaging was completed.

Brightfield images were acquired using an Olympus SZX16 stereomicroscope with either an Olympus DP26 or UC90 camera.

### Transmission Electron Microscopy

24 hpf embryos were fixed, stained and embedded using the microwave-assisted tissue processing protocol described in (Czopka and Lyons, 2011). Tails were dissected from embryos prior to fixation and used for genotyping.

Our tissue sampling and analytical techniques have been described previously in detail (Anderson et al., 2011a, 2011b; Lauritzen et al., 2013; Marc et al., 2013, 2014).

The tissues were osmicated for 60 min in 0.5% OsO4 in 0.1 M cacodylate buffer, processed in maleate buffer for en bloc staining with uranyl acetate, and processed for resin embedding. The epoxy resin bloc with zebrafish tissue was sectioned in the horizontal plane at 70–90 nm onto polyvinyl formal resin coated copper slot grids for transmission electron microscopy (TEM) (Lauritzen et al., 2013; Marc and Jones, 2002).

Each TEM section was imaged on a JEOL JEM-1400 transmission electron microscope at 20,000x and stored in 16- and 8-bit versions, as well as image pyramids of optimized tiles for web visualization with the Viking viewer (Anderson et al., 2011a, 2011b). Each image was captured as an array of image tiles at roughly 500-800 tiles/slice with 15% overlap.

### Image processing and analysis

Images were processed using ImageJ. Volume rendering was performed using FluoRender (Wan et al., 2009, 2017). For lateral view 3D rendering of the optic cup, the ectoderm was digitally erased in ImageJ prior to visualization in FluoRender. Invagination angles were measured as previously described (Bryan et al., 2016) and shown in Fig 2G. Individual cell tracking was performed as described in (Kwan et al., 2012) using LongTracker; nuclei were visualized using H2A.F/Z-mCherry.

## Acknowledgements

We are grateful to Rodney Stewart, David Grunwald, and Mick Jurynec for reagents, and the University of Utah Centralized Zebrafish Animal Resource for zebrafish husbandry. Thanks to members of the Kwan lab for useful discussions and critical reading of the manuscript. This work was supported by grants from the NEI/NIH to K. M. K. (R01 EY025378, R01 EY025780), to B. W. J. (R01 EY015128), and to the Moran Eye Center Vision Core (P30 EY014800). C. D. B. was supported by the University of Utah Developmental Biology Training Grant (NIH T32HD007491).

---

**Figure S1.**
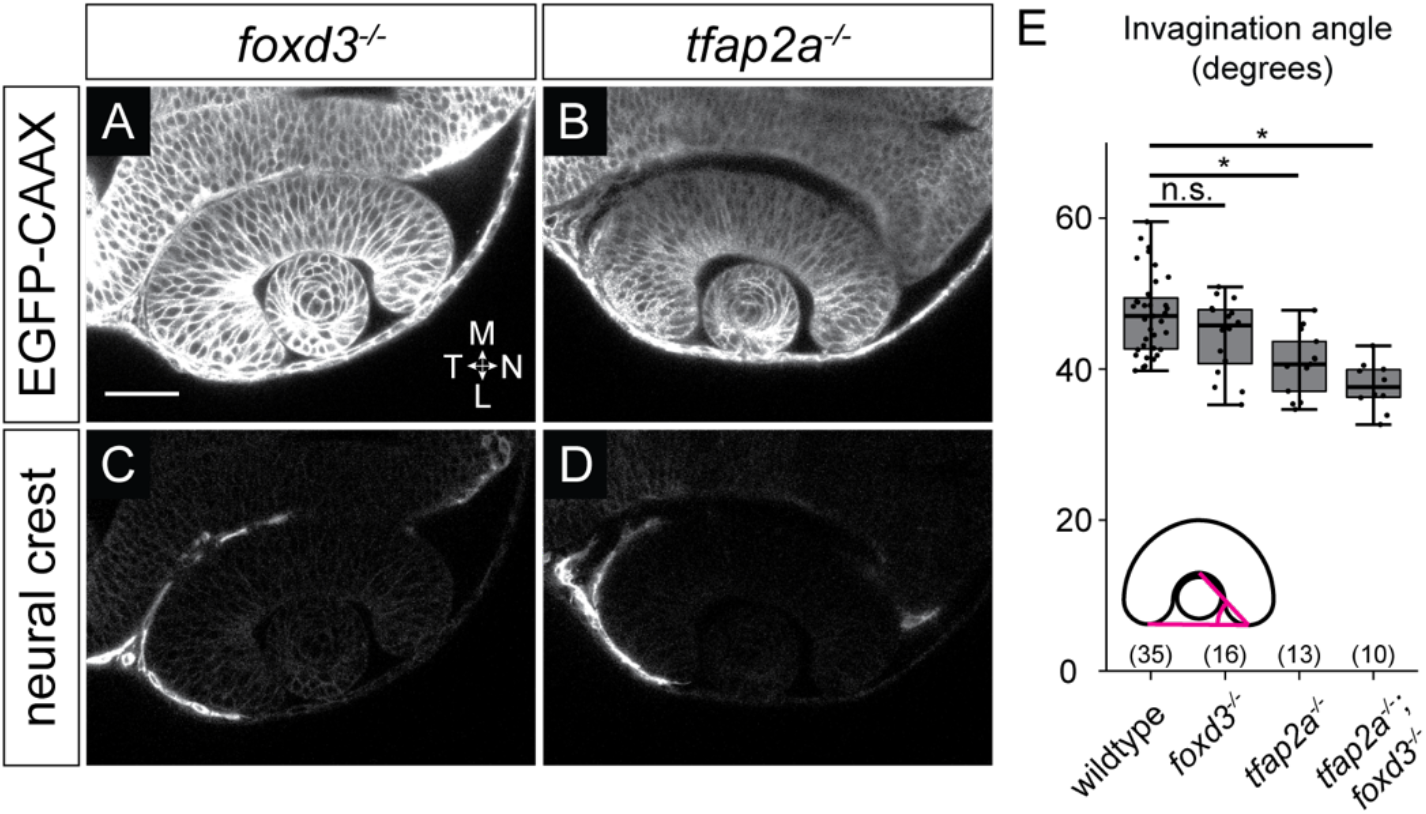
Invagination is disrupted in *tfap2a* but not *foxd3* single mutants. (A-D) Dorsal view, single confocal sections of 24 hpf *Tg(bactin2:EGFP-CAAX);Tg(sox10:memRFP)* double transgenic *foxd3* (A, C) and *tfap2a* (B, D) mutant embryos. Sections shown are at the dorsal/ventral midpoint of the lens. (E) Quantification of invagination angles, measured as shown in the inset diagram. ^*^P<0.001 using Welch’s t-test. (Wildtype vs. *tfap2a;foxd3* double mutant, P=2.52×10^-7^; wildtype vs. *tfap2a* single mutant, P=0.0001; wildtype vs. *foxd3* single mutant, P=0.07.) Results are from 3 experiments; n (embryos) shown in the base of the graph. Scale bar, 50 μm. M, medial; L, lateral; N, nasal; T, temporal.

**Figure S2.**
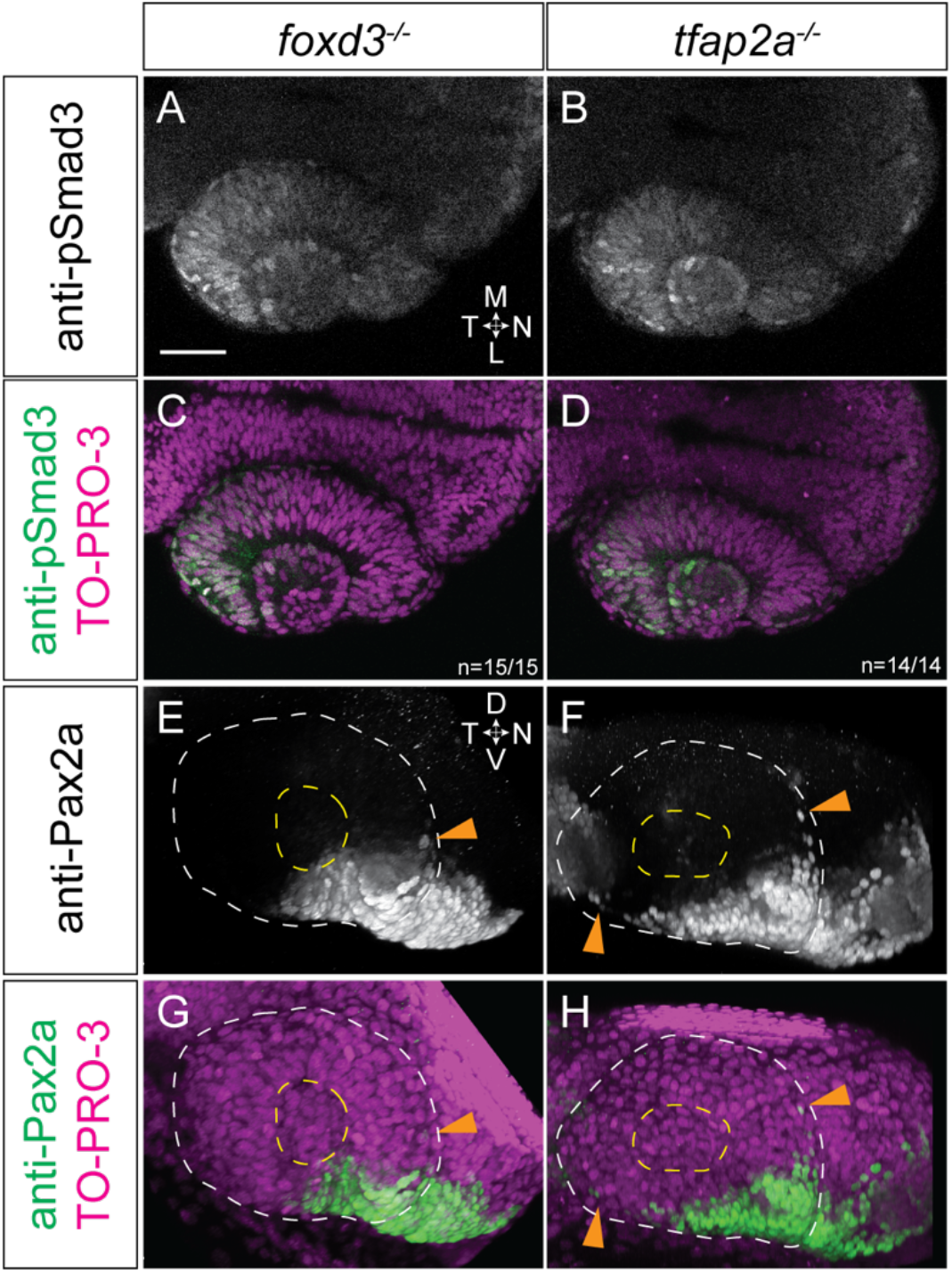
At 24 hpf, *tfap2a* and *foxd3* single mutants display normal TGF-beta signaling, while Pax2a expression expands into the RPE. (A-D) Dorsal view, single confocal sections of 24 hpf *foxd3* mutant (A, C) and *tfap2a* mutant (B,D) optic cups stained with anti-phospho-Smad3. Sections shown are at the dorsal/ventral lens midpoint. (E-H) Lateral view, 3D renderings of 24 hpf *foxd3* mutant (E, G) and *tfap2a* mutant (F,H) optic cups stained with anti-Pax2a. White dashed circles denote the boundary of the optic cup, yellow dashed circles display the boundary of the lens. Orange arrowheads in (E-H) indicate RPE cells which ectopically express Pax2a. Nuclei were counterstained with TO-PRO-3 (magenta); merges shown in (C, D, G, H). M, medial; L, lateral; D, dorsal; V, ventral; N, nasal; T, temporal.

**Figure S3.**
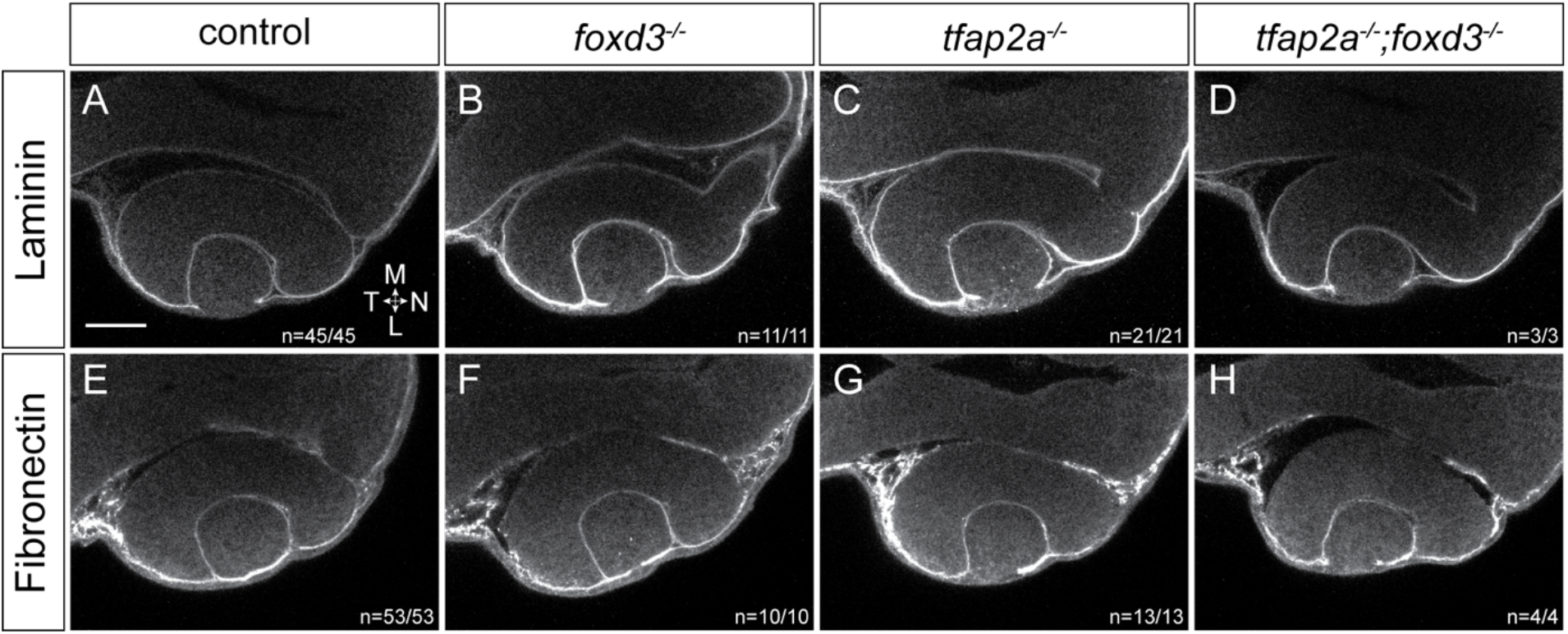
Laminin and fibronectin localization are unaffected in *tfap2a;foxd3* double mutants. At 24 hpf, both laminin (A-D) and fibronectin (E-H) are found around the developing optic cup in all genotypes shown. Dorsal view, single confocal sections. Scale bar, 50 μm. M, medial; L, lateral; N, nasal; T, temporal.

**Figure S4.**
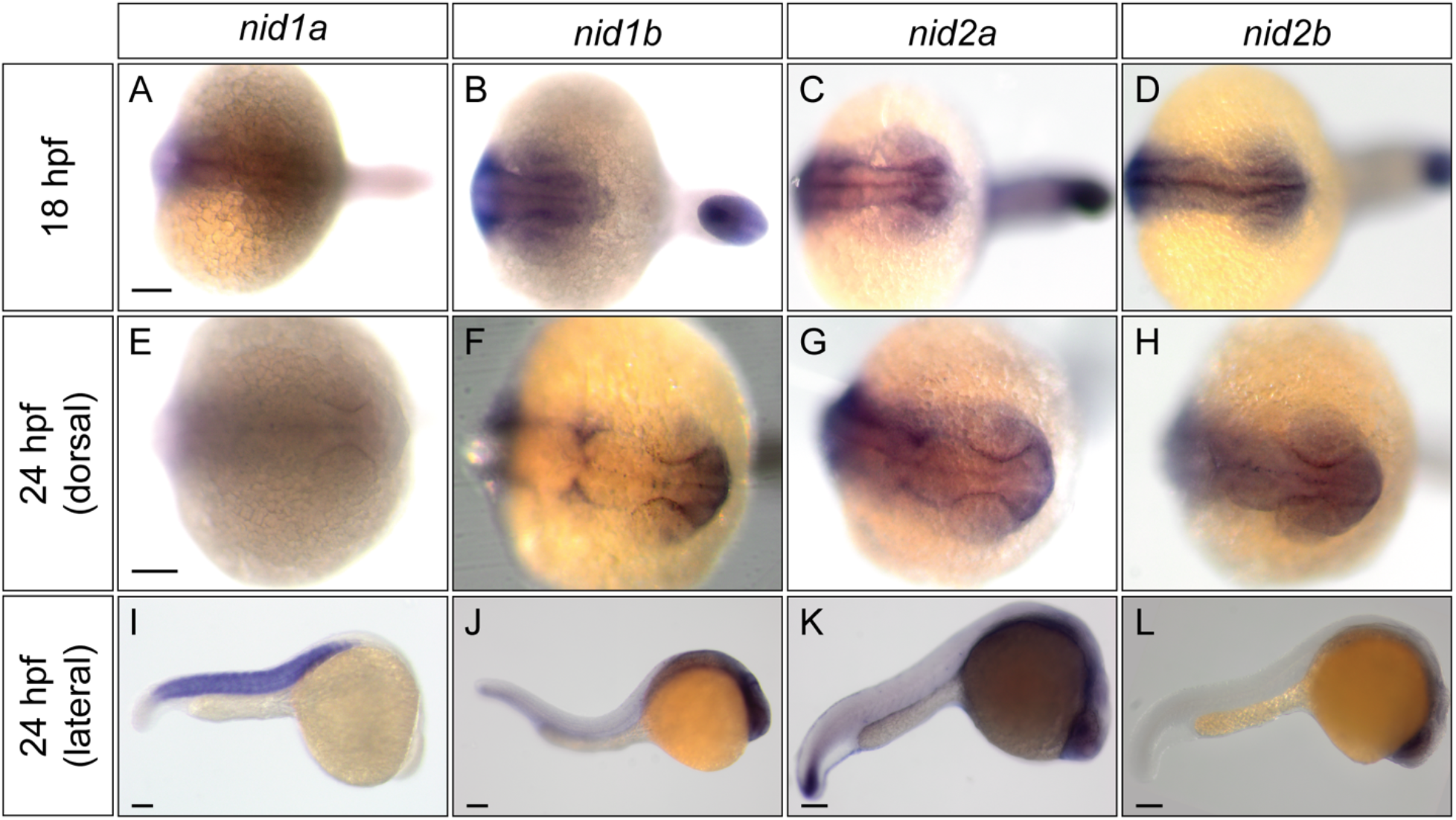
Zebrafish nidogen mRNA expression patterns at 18 and 24 hpf. Scale bars, 100 μm.

**Supplemental Movie 1.** Neural crest migration during optic cup morphogenesis. 12.5-24.5 hpf timelapse of a *Tg(β-actin2:EGFP-CAAX);Tg(sox10:memRFP)* double transgenic embryo. EGFP-CAAX labels all cell membranes (green), while membrane-bound RFP (magenta) labels only the neural crest. Dorsal view, single confocal section through the dorsal/ventral midpoint of the optic cup. Nasal (anterior) is to the right, temporal (posterior) to the left. ΔT between z-stacks, 3 minutes 30 seconds.

**Supplemental Movie 2.** Neural crest migration during optic cup morphogenesis. 12.5-24.5 hpf timelapse of a *Tg(β-actin2:EGFP-CAAX);Tg(sox10:memRFP)* double transgenic embryo. Lateral view, 3D rendering of the same timelapse dataset shown in Supplemental Movie 1. Only the RFP channel is shown (grayscale) to enable visualization of neural crest cell migration. Nasal (anterior) is to the right, temporal (posterior) to the left, dorsal to the top, ventral to the bottom. ΔT between z-stacks, 3 minutes 30 seconds

**Supplemental Movie 3.** Nuclear trajectories visualized in in three dimensions. 3D rendered rotation of 24 hpf timepoints showing representative trajectories of nuclei from cells in the nasal retina (orange), nasal RPE (blue) and temporal retina (green). Membrane channel is displayed in grayscale.

**Supplemental Movie 4.** Wildtype nasal RPE nuclear trajectories from 12.5-24 hpf. Representative trajectories over membrane channel average (grayscale). ΔT between z-stacks, 2 minutes 45 seconds.

**Supplemental Movie 5.** *tfap2a;foxd3* double mutant nasal RPE nuclear trajectories from 12.5-24 hpf. Representative trajectories over membrane channel average (grayscale). ΔT between z-stacks, 2 minutes 30 seconds.

**Supplemental Movie 6.** Wildtype nasal retina nuclear trajectories from 12.5-24 hpf. Representative trajectories over membrane channel average (grayscale). ΔT between z-stacks, 2 minutes 45 seconds.

**Supplemental Movie 7.** *tfap2a;foxd3* double mutant nasal retina nuclear trajectories from 12.5-24 hpf. Representative trajectories over membrane channel average (grayscale). ΔT between z-stacks, 2 minutes 30 seconds.

**Supplemental Movie 8.** Wildtype temporal retina nuclear trajectories from 12.5-24 hpf. Representative trajectories over membrane channel average (grayscale). ΔT between z-stacks, 2 minutes 45 seconds.

**Supplemental Movie 9.** *tfap2a;foxd3* double mutant temporal retina nuclear trajectories from 12.5-24 hpf. Representative trajectories over membrane channel average (grayscale). ΔT between z-stacks, 2 minutes 30 seconds.

## References

Akanuma, T., Koshida, S., Kawamura, A., Kishimoto, Y., and Takada, S. (2007). Paf1 complex homologues are required for Notch-regulated transcription during somite segmentation. EMBO Rep. 8, 858–863.

Anderson, J.R., Jones, B.W., Watt, C.B., Shaw, M. V, Yang, J.H., Demill, D., Lauritzen, J.S., Lin, Y., Rapp, K.D., Mastronarde, D., et al. (2011a). Exploring the retinal connectome. Mol Vis 17, 355–379.

Anderson, J.R., Mohammed, S., Grimm, B., Jones, B.W., Koshevoy, P., Tasdizen, T., Whitaker, R., and Marc, R.E. (2011b). The Viking viewer for connectomics: Scalable multi-user annotation and summarization of large volume data sets. J. Microsc. 241, 13–28.

Arduini, B.L., Bosse, K.M., and Henion, P.D. (2009). Genetic ablation of neural crest cell diversification. Development 136, 1987–1994.

Bader, B.L., Smyth, N., Nedbal, S., Baranowsky, A., Mokkapati, S., Miosge, N., Murshed, M., Nischt, R., Baranowsky, A., Mokkapati, S., et al. (2005). Compound Genetic Ablation of Nidogen 1 and 2 Causes Basement Membrane Defects and Perinatal Lethality in Mice. Mol. Cell. Biol. 25, 6846–6856.

Bassett, E.A., Williams, T., Zacharias, A.L., Gage, P.J., Fuhrmann, S., and West-Mays, J.A. (2010). AP-2a knockout mice exhibit optic cup patterning defects and failure of optic stalk morphogenesis. Hum. Mol. Genet. 19, 1791–1804.

Bohnsack, B.L., Kasprick, D.S., Kish, P.E., Goldman, D., and Kahana, A. (2012). A zebrafish model of Axenfeld-Rieger syndrome reveals that pitx2 regulation by Retinoic Acid is essential for ocular and craniofacial development. Investig. Ophthalmol. Vis. Sci. 53, 7–22.

Böse, K., Nischt, R., Page, A., Bader, B.L., Paulsson, M., and Smyth, N. (2006). Loss of nidogen-1 and -2 results in syndactyly and changes in limb development. J. Biol. Chem. 281, 39620–39629.

Bryan, C.D., Chien, C. Bin, and Kwan, K.M. (2016). Loss of laminin alpha 1 results in multiple structural defects and divergent effects on adhesion during vertebrate optic cup morphogenesis. Dev. Biol. 416, 324–337.

Chakraborty, S., Njah, K., Pobbati, A. V., Lim, Y.B., Raju, A., Lakshmanan, M., Tergaonkar, V., Lim, C.T., and Hong, W. (2017). Agrin as a Mechanotransduction Signal Regulating YAP through the Hippo Pathway. Cell Rep. 18, 2464–2479.

Cretekos, C.J., and Grunwald, D.J. (1999). Alyron, an Insertional Mutation Affecting Early Neural Crest Development in Zebrafish. Dev. Biol. 210, 322–338.

Czopka, T., and Lyons, D.A. (2011). Dissecting Mechanisms of Myelinated Axon Formation Using Zebrafish. In The Zebrafish: Disease Models and Chemical Screens, (Elsevier Inc.), pp. 25–62.

Das, A., and Crump, J.G. (2012). Bmps and Id2a act upstream of twist1 to restrict ectomesenchyme potential of the cranial neural crest. PLoS Genet. 8.

Dong, L.J., and Chung, A.E. (1991). The Expression of the Genes for Entactin, Laminin-a, Laminin-B1 and Laminin-B2 in Murine Lens Morphogenesis and Eye Development. Differentiation 48, 157–172.

Dutton, J.R., Antonellis, A., Carney, T.J., Rodrigues, F.S., Pavan, W.J., Ward, A., and Kelsh, R.N. (2008). An evolutionarily conserved intronic region controls the spatiotemporal expression of the transcription factor Sox10. BMC Dev. Biol. 8, 105.

Eberhart, J.K., He, X., Swartz, M.E., Yan, Y.L., Song, H., Boling, T.C., Kunerth, A.K., Walker, M.B., Kimmel, C.B., and Postlethwait, J.H. (2008). MicroRNA Mirn140 modulates Pdgf signaling during palatogenesis. Nat. Genet. 40, 290–298.

Eiraku, M., Takata, N., Ishibashi, H., Kawada, M., Sakakura, E., Okuda, S., Sekiguchi, K., Adachi, T., and Sasai, Y. (2011). Self-organizing optic-cup morphogenesis in three-dimensional culture. Nature 472, 51–56.

Ekblom, P., Ekblom, M., Fecker, L., Klein, G., Zhang, H.Y., Kadoya, Y., Chu, M.L., Mayer, U., and Timpl, R. (1994). Role of mesenchymal nidogen for epithelial morphogenesis in vitro. Development 120, 2003–2014.

Fox, J.W., Mayer, U., Nischt, R., Aumailley, M., Reinhardt, D., Wiedemann, H., Mann, K., Timpl, R., Krieg, T., and Engel, J. (1991). Recombinant nidogen consists of three globular domains and mediates binding of laminin to collagen type IV. EMBO J. 10, 3137–3146.

Fuhrmann, S., Levine, E.M., and Reh, T.A. (2000). Extraocular mesenchyme patterns the optic vesicle during early eye development in the embryonic chick. Development 127, 4599–4609.

Gage, P.J., Rhoades, W., Prucka, S.K., and Hjalt, T. (2005). Fate maps of neural crest and mesoderm in the mammalian eye. Investig. Ophthalmol. Vis. Sci. 46, 4200–4208.

Gestri, G., Osborne, R.J., Wyatt, A.W., Gerrelli, D., Gribble, S., Stewart, H., Fryer, A., Bunyan, D.J., Prescott, K., Collin, J.R.O., et al. (2009). Reduced TFAP2A function causes variable optic fissure closure and retinal defects and sensitizes eye development to mutations in other morphogenetic regulators. Hum. Genet. 126, 791–803.

Gestri, G., Bazin-Lopez, N., Scholes, C., and Wilson, S.W. (2018). Cell Behaviors during Closure of the Choroid Fissure in the Developing Eye. Front. Cell. Neurosci. 12, 1–12.

Grocott, T., Johnson, S., Bailey, A.P., and Streit, A. (2011). Neural crest cells organize the eye via TGF-β and canonical Wnt signalling. Nat. Commun. 2, 266–269.

Hayes, J.M., Hartsock, A., Clark, B.S., Napier, H.R.L., Link, B.A., and Gross, J.M. (2012). Integrin 5/Fibronectin1 and focal adhesion kinase are required for lens fiber morphogenesis in zebrafish. Mol. Biol. Cell 23, 4725–4738.

Heermann, S., Schütz, L., Lemke, S., Krieglstein, K., and Wittbrodt, J. (2015). Eye morphogenesis driven by epithelial flow into the optic cup facilitated by modulation of bone morphogenetic protein. Elife 4, 1–17.

Hendrix, R.W., and Zwaan, J. (1975). The matrix of the optic vesicle-presumptive lens interface during induction of the lens in the chicken embryo. J. Embryol. Exp. Morphol. 33, 1023–1049.

Hero, I. (1990). Optic fissure closure in the normal cinnamon mouse: An ultrastructural study. Investig. Ophthalmol. Vis. Sci. 31, 197–216.

Hero, I., Farjah, M., and Scholtz, C.L. (1991). The prenatal development of the optic fissure in colobomatous microphthalmia. Investig. Ophthalmol. Vis. Sci. 32, 2622–2635.

Hilfer, S.R. (1983). Development of the eye of the chick embryo. Scan. Electron Microsc. 1353–1369.

Huang, J., Rajagopal, R., Liu, Y., Dattilo, L.K., Shaham, O., Ashery-Padan, R., and Beebe, D.C. (2011). The mechanism of lens placode formation: A case of matrix-mediated morphogenesis. Dev. Biol. 355, 32–42.

Ivanovitch, K., Cavodeassi, F., and Wilson, S.W. (2013). Precocious acquisition of neuroepithelial character in the eye field underlies the onset of eye morphogenesis. Dev. Cell 27, 293–305.

James, A., Lee, C., Williams, A.M., Angileri, K., Lathrop, K.L., and Gross, J.M. (2016). The hyaloid vasculature facilitates basement membrane breakdown during choroid fissure closure in the zebrafish eye. Dev. Biol. 419, 262–272.

Johnston, M.C., Noden, D.M., Hazelton, R.D., Coulombre, J.L., and Coulombre, A.J. (1979). Origins of avian ocular and periocular tissues. Exp. Eye Res. 29, 27–43.

Kadoya, Y., Salmivirta, K., Talts, J.F., Kadoya, K., Mayer, U., Timpl, R., and Ekblom, P. (1997). Importance of nidogen binding to laminin γ1 for branching epithelial morphogenesis of the submandibular gland. Development 124, 683–691.

Kheradmand, F., Rishi, K., and Werb, Z. (2002). Signaling through the EGF receptor controls lung morphogenesis in part by regulating MT1-MMP-mediated activation of gelatinase A/MMP2. J. Cell Sci. 115, 839–848.

Kimmel, C.B., Ballard, W.W., Kimmel, S.R., Ullmann, B., and Schilling, T.F. (1995). Stages of embryonic development of the zebrafish. Dev. Dyn. 203, 253–310.

Kirby, B.B., Takada, N., Latimer, A.J., Shin, J., Carney, T.J., Kelsh, R.N., and Appel, B. (2006). In vivo time-lapse imaging shows dynamic oligodendrocyte progenitor behavior during zebrafish development. Nat. Neurosci. 9, 1506–1511.

Knight, R.D., Javidan, Y., Zhang, T., Nelson, S., and Schilling, T.F. (2005). AP2-dependent signals from the ectoderm regulate craniofacial development in the zebrafish embryo. Development 132, 3127–3138.

Kudoh, T., Tsang, M., Hukriede, N.A., Chen, X., Dedekian, M., Clarke, C.J., Kiang, A., Schultz, S., Epstein, J.A., Toyama, R., et al. (2001). A gene expression screen in zebrafish embryogenesis. Genome Res. 11, 1979–1987.

Kwan, K.M. (2014). Coming into focus: The role of extracellular matrix in vertebrate optic cup morphogenesis. Dev. Dyn. 243, 1242–1248.

Kwan, K.M., Fujimoto, E., Grabher, C., Mangum, B.D., Hardy, M.E., Campbell, D.S., Parant, J.M., Yost, H.J., Kanki, J.P., and Chien, C. Bin (2007). The Tol2kit: A multisite gateway-based construction Kit for Tol2 transposon transgenesis constructs. Dev. Dyn. 236, 3088–3099.

Kwan, K.M., Otsuna, H., Kidokoro, H., Carney, K.R., Saijoh, Y., and Chien, C.-B. (2012). A complex choreography of cell movements shapes the vertebrate eye. Development 139, 359–372.

Langenbacher, A.D., Nguyen, C.T., Cavanaugh, A.M., Huang, J., Lu, F., and Chen, J.N. (2011). The PAF1 complex differentially regulates cardiomyocyte specification. Dev. Biol. 353, 19–28.

Langenberg, T., Kahana, A., Wszalek, J.A., and Halloran, M.C. (2008). The eye organizes neural crest cell migration. Dev. Dyn. 237, 1645–1652.

Lauritzen, J.S., Anderson, J.R., Jones, B.W., Watt, C.B., Mohammed, S., Hoang, J. V., and Marc, R.E. (2013). ON Cone Bipolar Cell Axonal Synapses in the OFF Inner Plexiform Layer of the Rabbit Retina. J. Comp. Neurol. 521, 977–1000.

Lauter, G., Söll, I., and Hauptmann, G. (2014). Sensitive Whole-Mount Fluorescent In Situ Hybridization in Zebrafish Using Enhanced Tyramide Signal Amplification. Methods Mol. Biol. 1082, 175–185.

Lee, J., Willer, J.R., Willer, G.B., Smith, K., Gregg, R.G., and Gross, J.M. (2008). Zebrafish blowout provides genetic evidence for Patched1-mediated negative regulation of Hedgehog signaling within the proximal optic vesicle of the vertebrate eye. Dev. Biol. 319, 10–22.

Leerberg, D.M., Sano, K., and Draper, B.W. (2017). Fibroblast growth factor signaling is required for early somatic gonad development in zebrafish. PLOS Genet. 1–28.

Li, W., and Cornell, R.A. (2007). Redundant activities of Tfap2a and Tfap2c are required for neural crest induction and development of other non-neural ectoderm derivatives in zebrafish embryos. Dev. Biol. 304, 338–354.

Li, Z., Joseph, N.M., and Easter, S.S. (2000). The morphogenesis of the zebrafish eye, including a fate map of the optic vesicle. Dev. Dyn. 218, 175–188.

Lupo, G., Gestri, G., O’Brien, M., Denton, R.M., Chandraratna, R.A.S., Ley, S. V, Harris, W.A., and Wilson, S.W. (2011). Retinoic acid receptor signaling regulates choroid fissure closure through independent mechanisms in the ventral optic cup and periocular mesenchyme. Proc. Natl. Acad. Sci. U. S. A. 108, 8698–8703.

Marc, R.E., and Jones, B.W. (2002). Molecular phenotyping of retinal ganglion cells. J. Neurosci. 22, 413–427.

Marc, R.E., Jones, B.W., Watt, C.B., Anderson, J.R., Sigulinsky, C., and Lauritzen, S. (2013). Retinal connectomics: Towards complete, accurate networks. Prog. Retin. Eye Res. 37, 141–162.

Marc, R.E., Anderson, J.R., Jones, B.W., Sigulinsky, C.L., and Lauritzen, J.S. (2014). The AII amacrine cell connectome: a dense network hub. Front. Neural Circuits 8, 1–13.

Martinez-Morales, J.R., Rembold, M., Greger, K., Simpson, J.C., Brown, K.E., Quiring, R., Pepperkok, R., Martin-Bermudo, M.D., Himmelbauer, H., and Wittbrodt, J. (2009). Ojoplano-mediated basal constriction is essential for optic cup morphogenesis. Development 136, 2165–2175.

Miesfeld, J.B., Gestri, G., Clark, B.S., Flinn, M.A., Poole, R.J., Bader, J.R., Besharse, J.C., Wilson, S.W., and Link, B.A. (2015). Yap and Taz regulate retinal pigment epithelial cell fate. Development 142, 3021–3032.

Neff, M.M., Neff, J.D., Chory, J., and Pepper, A.E. (1998). dCAPS, a simple technique for the genetic analysis of single nucleotide polymorphisms: experimental applications in Arabidopsis thaliana genetics. Plant J. 14, 387–392.

Nelson, D.A., and Larsen, M. (2015). Heterotypic control of basement membrane dynamics during branching morphogenesis. Dev. Biol. 401, 103–109.

Nguyen, C.T., Langenbacher, A., Hsieh, M., and Chen, J.N.O. (2010). The PAF1 complex component Leo1 is essential for cardiac and neural crest development in zebrafish. Dev. Biol. 341, 167–175.

Nicolás-Pérez, M., Kuchling, F., Letelier, J., Polvillo, R., Wittbrodt, J., and Martínez-Morales, J.R. (2016). Analysis of cellular behavior and cytoskeletal dynamics reveal a constriction mechanism driving optic cup morphogenesis. Elife 5, 1–24.

Peterson, P.E., Pow, C.S., Wilson, D.B., and Hendrickx, A.G. (1995). Localisation of glycoproteins and glycosaminoglycans during early eye development in the macaque. J. Anat. 186 ( Pt 1, 31–42.

Picker, A., Cavodeassi, F., Machate, A., Bernauer, S., Hans, S., Abe, G., Kawakami, K., Wilson, S.W., and Brand, M. (2009). Dynamic coupling of pattern formation and morphogenesis in the developing vertebrate retina. PLoS Biol. 7.

Provost, E., Rhee, J., and Leach, S.D. (2007). Viral 2A peptides allow expression of multiple proteins from a single ORF in transgenic zebrafish embryos. Genesis 45, 625–629.

Pujuguet, P., Simian, M., Liaw, J., Timpl, R., Werb, Z., and Bissell, M.J. (2000). Nidogen-1 regulates laminin-1-dependent mammary-specific gene expression. J. Cell Sci. 113, 849–858.

Reinhardt, D., Mann, K., Nischt, R., Fox, J.W., Chu, M.L., Krieg, T., and Timpl, R. (1993). Mapping of nidogen binding sites for collagen type IV, heparan sulfate proteoglycan, and zinc. J. Biol. Chem. 268, 10881–10887.

Schmitt, E.A., and Dowling, J.E. (1994). Early eye morphogenesis in the zebrafish, Brachydanio rerio. J. Comp. Neurol. 344, 532–542.

Schook, P. (1980). Morphogenetic movements during the early development of the chick eye. A light microscopic and spatial reconstructive study. Acta Morphol. Neerl. Scand. 18, 1–30.

Sedykh, I., Yoon, B., Roberson, L., Moskvin, O., Dewey, C.N., and Grinblat, Y. (2017). Zebrafish zic2 controls formation of periocular neural crest and choroid fissure morphogenesis. Dev. Biol. 429, 92–104.

Senior, R.M., Griffin, G.L., Mudd, M.S., Moxley, M.A., Longmore, W.J., and Pierce, R.A. (1996). Entactin expression by rat lung and rat alveolar epithelial cells. Am. J. Respir. Cell Mol. Biol. 14, 239–247.

Sidhaye, J., and Norden, C. (2017). Concerted action of neuroepithelial basal shrinkage and active epithelial migration ensures efficient optic cup morphogenesis. Elife 6, 1–29.

Soules, K.A., and Link, B.A. (2005). Morphogenesis of the anterior segment in the zebrafish eye. BMC Dev. Biol. 5, 12.

Sowden, J.C. (2007). Molecular and developmental mechanisms of anterior segment dysgenesis. Eye (Lond). 21, 1310–1318.

Svoboda, K.K., and O’Shea, K.S. (1987). An analysis of cell shape and the neuroepithelial basal lamina during optic vesicle formation in the mouse embryo. Development 100, 185–200.

Thesleff, I. (2003). Epithelial-mesenchymal signalling regulating tooth morphogenesis. J. Cell Sci. 116, 1647–1648.

Thisse, B., and Thisse, C. (2004). Fast Release Clones: A High Throughput Expression Analysis. ZFIN Direct Data Submiss.

Thisse, C., and Thisse, B. (2008). High-resolution in situ hybridization to whole-mount zebrafish embryos. Nat. Protoc. 3, 59–69.

Tuckett, F., and Morriss-Kay, G.M. (1986). The distribution of fibronectin, laminin and entactin in the neurulating rat embryo studied by indirect immunofluorescence. J. Embryol. Exp. Morphol. 94, 95–112.

Walls, G.L. (1942). The vertebrate eye and its adaptive radiation (Hafner Publishing Company).

Wan, Y., Otsuna, H., Chien, C. Bin, and Hansen, C. (2009). An interactive visualization tool for multi-channel confocal microscopy data in neurobiology research. IEEE Trans. Vis. Comput. Graph. 15, 1489–1496.

Wan, Y., Otsuna, H., Holman, H.A., Bagley, B., Ito, M., Lewis, A.K., Colasanto, M., Kardon, G., Ito, K., and Hansen, C. (2017). FluoRender: Joint freehand segmentation and visualization for many-channel fluorescence data analysis. BMC Bioinformatics 18, 1–15.

Wang, W.-D., Melville, D.B., Montero-Balaguer, M., Hatzopoulos, A.K., and Knapik, E.W. (2011). Tfap2a and Foxd3 regulate early steps in the development of the neural crest progenitor population. Dev. Biol. 360, 173–185.

Weiss, O., Kaufman, R., Michaeli, N., and Inbal, A. (2012). Abnormal vasculature interferes with optic fissure closure in lmo2 mutant zebrafish embryos. Dev. Biol. 369, 191–198.

Wells, K.L., Gaete, M., Matalova, E., Deutsch, D., Rice, D., and Tucker, A.S. (2013). Dynamic relationship of the epithelium and mesenchyme during salivary gland initiation: the role of Fgf10. Biol. Open 2, 981–989.

Willem, M., Miosge, N., Halfter, W., Smyth, N., Jannetti, I., Burghart, E., Timpl, R., and Mayer, U. (2002). Specific ablation of the nidogen-binding site in the laminin γ1 chain interferes with kidney and lung development. Development 129, 2711–2722.

Williams, A.L., and Bohnsack, B.L. (2015). Neural crest derivatives in ocular development: Discerning the eye of the storm. Birth Defects Res. Part C - Embryo Today Rev. 105, 87–95.

Zhu, P., Ma, Z., Guo, L., Zhang, W., Zhang, Q., Zhao, T., Jiang, K., Peng, J., and Chen, J. (2017). Short body length phenotype is compensated by the upregulation of nidogen family members in a deleterious nid1a mutation of zebrafish. J. Genet. Genomics 44, 553–556.

